# A single-cell, time-resolved profiling of Xenopus mucociliary epithelium reveals non-hierarchical model of development

**DOI:** 10.1101/2023.01.03.521555

**Authors:** Julie Lee, Andreas Fønss Møller, Shinhyeok Chae, Alexandra Bussek, Tae Joo Park, Youni Kim, Hyun-Shik Lee, Tune H. Pers, Taejoon Kwon, Jakub Sedzinski, Kedar Nath Natarajan

## Abstract

The specialized cell-types of the mucociliary epithelium (MCE) lining the respiratory tract enable continuous airway clearing, with its defects leading to chronic respiratory diseases. The molecular mechanisms driving cell-fate acquisition and temporal specialization during mucociliary epithelial development remain largely unknown. Here, we profile the developing *Xenopus* MCE from pluripotent to mature stages by single-cell transcriptomics, identifying novel, multipotent *early epithelial progenitors* that execute multi-lineage cues before specialising into late-stage ionocytes, goblet and basal cells. Combining *in silico* lineage inference, *in situ* hybridization and single-cell multiplexed RNA imaging, we capture the initial bifurcation into early epithelial and multiciliated progenitors, chart cell-type emergence and fate progression into specialized cell-types. Comparative analysis of nine airway atlases reveals an evolutionary conserved transcriptional module in ciliated cells, whereas secretory and basal types execute distinct function-specific programmes across vertebrates. We uncover a continuous non-hierarchical model of MCE development alongside a significant data resource for understanding respiratory biology.

## INTRODUCTION

Embryonic development is a tightly regulated process, where spatio-temporal organisational cues drive cell fate specification and the emerging complexity of cell-types within an embryo. The series of single-cell responses and collective cell state transitions guide the organism’s body plan and tissue organisation. Of fundamental importance is the development of the vertebrate airway and the lining of the mucociliary epithelium (MCE), which enables fundamental respiratory air exchange and critical defence against inhaled agents, microorganisms and harmful substances (*1–4*). During development, the mammalian MCE forms as a pseudostratified complex tissue consisting of multiple cell-types, further regulated by chemical and mechanical cues within the epithelium (*5, 6*). The outer epithelial mature goblet cells secrete mucus that along with anti-inflammatory peptides maintains a physical and chemical barrier between the respiratory epithelium and the inhaled air (*7*). The multiciliated cells and coordinated beating of their cilia propels mucus to the oropharynx where it is expectorated or swallowed (*8*). The basal cells located below the superficial epithelial cell layer are thought to differentiate into other specialised mucociliary cells across late stages (*9*), and the proportionally fewer ionocytes control mucus viscosity by balancing electrolyte homeostasis across the epithelium (*10*). These mature MCE cell-types structurally form the organism’s essential respiratory architecture, acting in concert as the primary innate airway defence barrier, and through efficient mucus clearance govern essential airway conductance and molecular transport. The increased susceptibility to airway infections, respiratory diseases and impaired lung function during asthma, chronic obstructive pulmonary disease, and cystic fibrosis is characterised by altered composition, abundance and distribution of airway MCE cell-types (*10–14*)

The embryonic epidermis of the amphibian *Xenopus* has emerged as a powerful model to study vertebrate mucociliary epithelium. Similar to the mammalian airway, the *Xenopus* epidermis develops as a mix of multiciliated and secretory cells (*15*). The *Xenopus* and mammalian mucociliary epithelia share striking similarities, with many protein counterparts recently discovered in mammalian airways (*16–19*). Recent studies have explored individual MCE cell-types and signalling pathways during MCE regeneration and homeostasis across many species (*20–23*). Applications of single-cell transcriptomics in *Xenopus* species across embryonic, larval and adult stages are enabling large data resources to study development (*24, 25*). However, the molecular mechanisms underlying cell-fate acquisition, cell-type compositions and temporal dynamics, particularly during MCE development have been lacking. The blastula stage *Xenopus* embryos (*animal cap*) composed of pluripotent cell sheets can be cultured as *ex vivo* explants (*organoids*), and undergo default mucociliary fate specification into a functional MCE robustly mimicking the earliest developmental events (*26–28*). Here, we profile the entire MCE development across ten stages through single-cell transcriptomics (scRNA-seq) using *Xenopus* organoids and perform the validation by *in situ* hybridization and multiplexed spatial RNA imaging in *Xenopus* embryos. Profiling 33,990 single-cells, we characterise the developmental diversification of transitory early-stage cell states and their specialisation into basal cells and mature multiciliated, goblet cells and ionocytes. Notably, we identify and capture a new multipotent ‘early epithelial progenitor’ population at *neurula* developmental stages. Devising a generalizable *in-silico* lineage inference method, we demonstrate that basal and secretory (*i.e.* goblet cell and ionocyte) differentiation requires passage through the multipotent early epithelial progenitors, while ciliated progenitors mature across a distinct lineage into multiciliated cells. We validate the timing, appearance, spatial positioning and specialisation of different cell-types through single-cell *in-situ* HCR and multiplexed RNA imaging. Through comparative analysis of nine cross-species single-cell airway atlases and corresponding cell-types, we identify a conserved species-independent expression programme in ciliated cells, alongside cell-type and species-specific functional programmes within basal and secretory cell-types (Fig. 1A). Collectively, our work provides a temporal atlas and associated dynamics of emerging specialised cell-types, laying the building foundation for dissecting the stepwise formation of complex mucociliary epithelium.

**Fig. 1.**
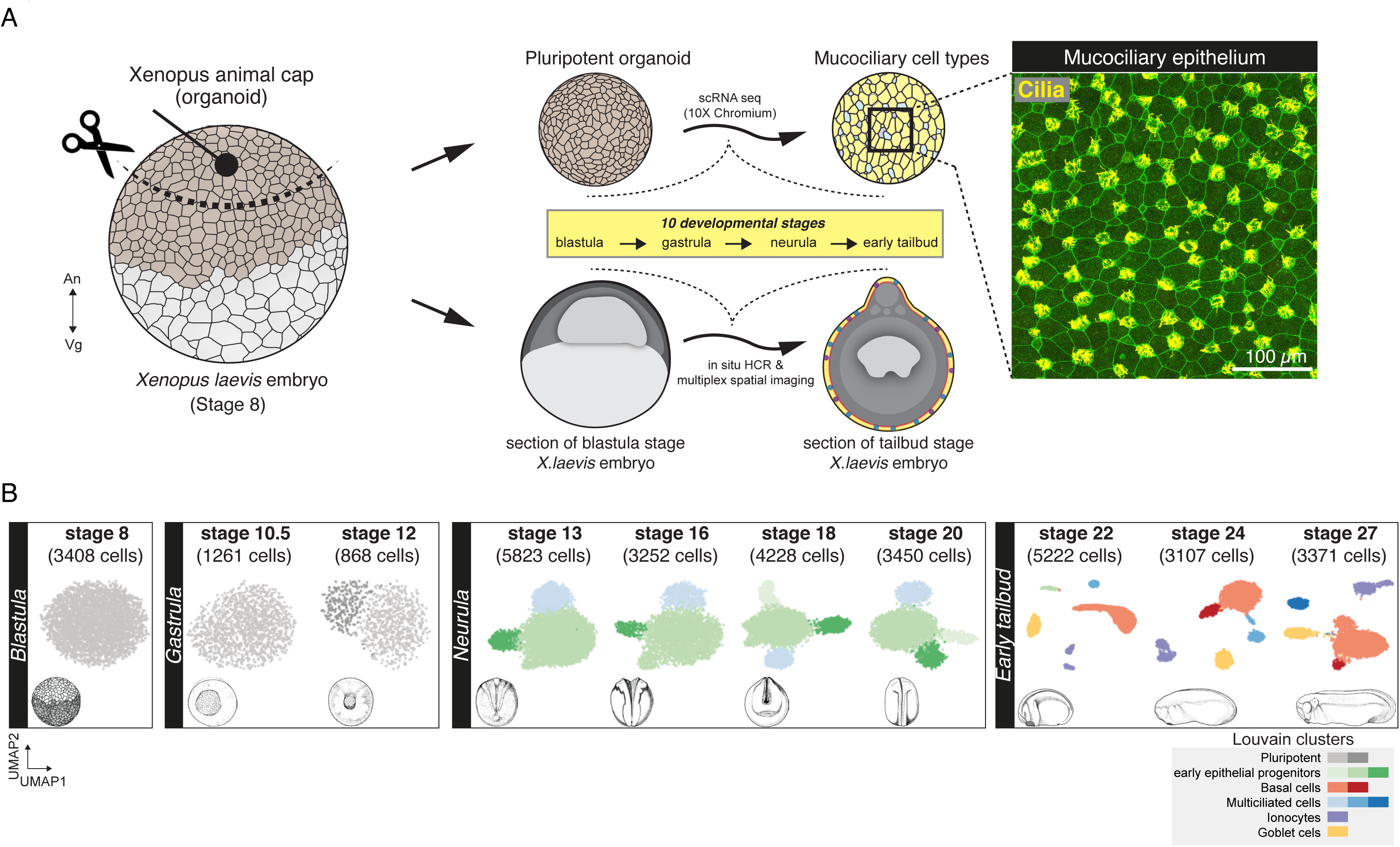
Cellular composition of developing *Xenopus* mucociliary epithelium (MCE) **(A)** Schema for single-cell transcriptome profiling of developing *Xenopus* MCE. The animal pole tissues from blastula stage 8 embryos were cultured as organoids, and sampled temporally across the 10 developmental stages (spanning blastula, gastrula, neurula and early tailbud stages) and profiled by droplet scRNA-seq. Inset: surface mucociliary epithelium of stage 27 organoids stained with anti-acetylated tubulin antibody (marking axonemes that build cilia, yellow) and phalloidin (marking filamentous actin, green). (B) Low dimensional (UMAP) scRNA-seq visualisation of the different developmental stages and Louvain clusters, colored based on major cell-types.

## RESULTS

### Single-cell transcriptomic profiling of developing mucociliary epithelium

To generate a continuous fate map of developing *Xenopus* MCE, we profiled animal cap organoids corresponding to ten developmental stages from early pluripotency to a functional MCE spanning *blastula* (Nieuwkoop and Faber (NF) stage 8), *gastrula* (stages 10.5 and 12), *neurula* (stages 13, 16, 18 and 21) and *early tailbud* development (stages 22, 24 and 27) (*29*) (Fig. 1A; methods); obtaining transcriptomes of 33,990 single-cells (Fig. S1A-C; methods, supplementary note 1). The initial assessment of individual stages by unsupervised clustering revealed the formation of all known major MCE cell-types at early tailbud stages and across several sub-clusters (Fig. 1B, S2A-C, Table S2) (*30, 31*).

At the blastula stage, the fate-unspecified pluripotent cells formed a homogeneous population expressing pluripotency transcription factors *pou5f3* and other markers of the undifferentiated state such as *sox3* (*32, 33*) (Fig. 1B, S2A), which could be further distinguished into two clusters by gastrula stages (dark and light grey clusters, respectively; Fig. 1B, S2A). The neurula stages (stages 13-20) were characterised by a complete loss of pluripotency and emergence of previously unknown progenitor populations (green and blue clusters, Fig. 1B, S2B). We termed the larger population as early epithelial progenitors (three subclusters), expressing stochastic, low expression levels of multiple genes such as extracellular matrix component (*has1*), elongation factor protein (*eef1a1o*) at single-cell level (Fig. 1B, S2B). Intriguingly, by late neurula, the individual early epithelial progenitors subclusters expressed secretory markers (goblet: glycoprotein otogelin *(otog,* also known as *mucXS* and *otogl2)* and ionocytes: vacuolar-type ATPase proton transporter (*atp6v1g3*), indicating their maturation into goblet cells and ionocytes (Fig. S2B). The smaller neurula stage population (light blue; Fig. S2B) expressed microtubule components (beta-tubulin: *tubb4b*, dynein: *dynll1*), indicating a bias towards multiciliated cells. The tailbud stages (stages 22-27) marked the respective specialisation of the early multiciliated population into multiciliated clusters, and early epithelial progenitor subpopulations into ionocytes, goblet, and basal cell clusters (Fig. 1B, S2C). The initial stage-specific scRNA-seq analysis captured major MCE cell-types but with previously unknown heterogeneity within cell-types.

### Integrated time-course analysis of mucociliary epithelial development

To capture the developmental progression and single-cell state transitions over the entire MCE development, we integrated, visualised, and analysed all the stages together over a graph embedding of the developmental manifold (Fig. 2A, Table S5, visualization portal). This low-dimensional embedding mimicked a starfish pattern, with the blastula, and gastrula stages at the head (stages 8, 10.5, 12) separated from the late stages across individual arms (stages 22-27). Graph-based approaches are suited for high-density datasets, and conventional community-based clustering approaches failed to capture meaningful cell states and types (Fig. S3D). To perform classification over the continuous high cell density MCE developmental manifold, we adapted and applied Phenograph clustering (*34*), and identified five major cell-types, separated into 15 different clusters (Fig. 2B, S3A-B, Table S3-S6). These clusters included two pluripotent (P1 and P2), four early epithelial progenitors (Eep1, Eep2, Eep3 and Eep4), four basal (Bc1, Bc2, Bc3 and Bc4), three multiciliated (Mcc1, Mcc2 and Mcc3), an ionocyte (Ic) and goblet cell clusters (Gc) (Fig. 2A-B & S3A-B). The clusters were named based on the expression of known and additional new cell-type specific markers (Fig. 2C-D, S3C, S4A-F). Notably, the cluster membership spanned single-cells from multiple developmental stages and scRNA-seq runs, indicating that clusters are biologically relevant and not due to technical or batch effects (Fig. 2A-D, S3C, S4A-F). While the major cell-types are discrete, the cell states (i.e. sub-clusters) are transitory and describe the different trajectories taken by single-cells over the developmental MCE manifold.

**Fig. 2.**
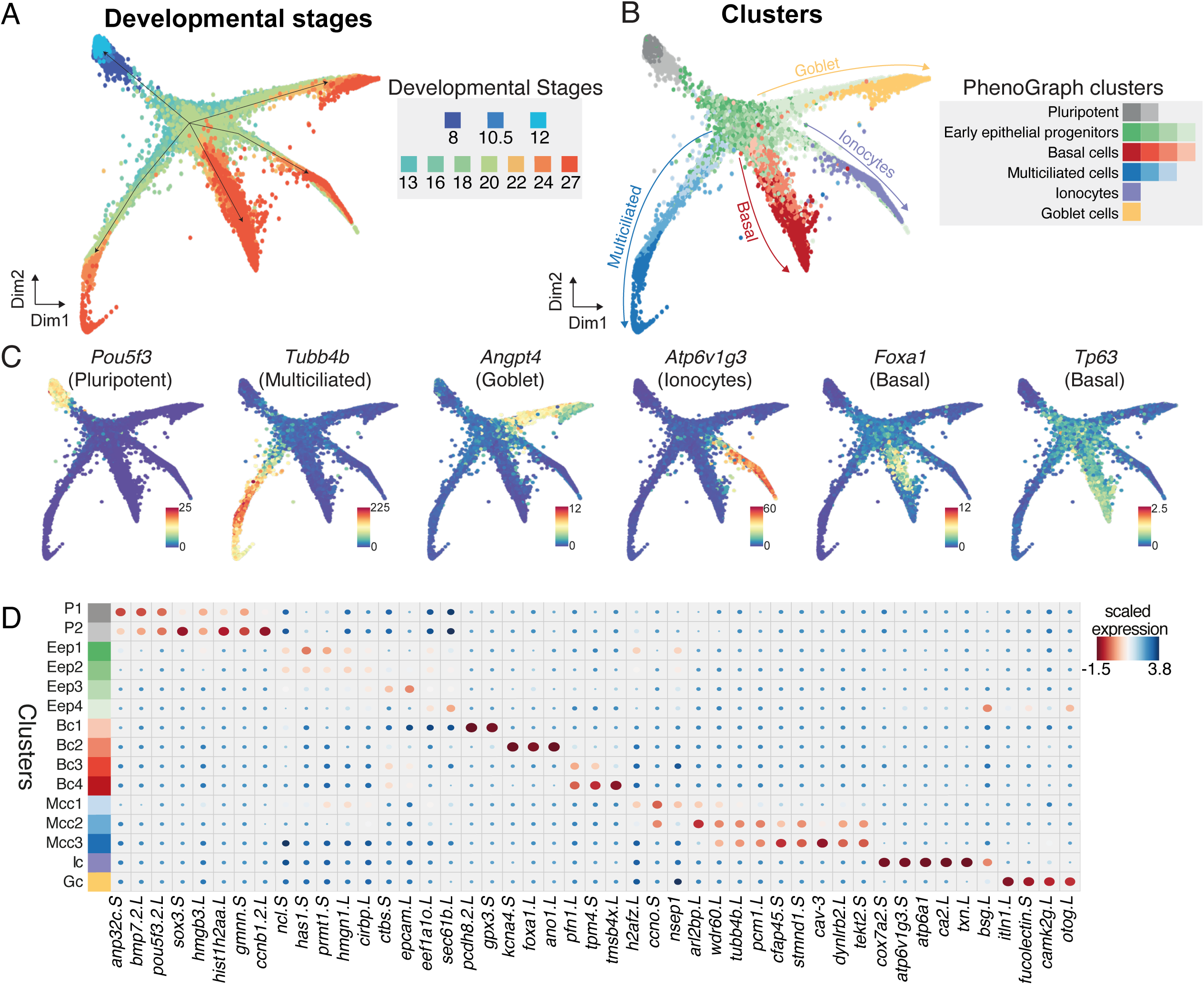
Joint embedding of continuous MCE developmental manifold and cell-type differentiation. **(A)** High-density force-directed k-nearest-neighbour (knn) graph visualisation of single-cells, colored by developmental stages. **(B)** PhenoGraph clustering of single-cells over continuous MCE manifold, colored by different cell-types and cell states (subclusters). The arrows indicate the differentiation of progenitors into specific cell-types. (C) Expression patterns of marker genes for MCE cell-types overlaid on the knn-graph. The scale bars indicated scaled imputed expression levels. (D) Dot plot of marker genes across different MCE clusters. The colour represents maximum-normalised mean marker gene expression across each cluster, and size indicates the proportion of positive cells relative to the entire dataset.

The pluripotent clusters (P1, P2) expressed respective transcription factors (TFs) (*pou5f3, sox3)*, mRNA splicing factors (*srsf5, srsf7*), signalling molecules (*bmp7.2*) and cell cycle regulators (*anp32c*, *gmnn*), indicating a hyper-transcriptional activity and fast cycling cells (Fig. 2C, S3C, S4A), consistent with the highest number of highly variable genes (Fig. S1C). The P1 and P2 clusters could be distinguished based on the expression of *krt5.7.L*, *upk3b.L* and *grhl3.S*, which become restricted to cells present in the outer or superficial layers (*35*). The early epithelial progenitors (Eep1-4) expressed several general TFs (*nsep1*, *prmt1*, *hmgn1*) as well as genes with broad cellular functions including adhesion (*epcam*), poly-A binding (*ncl*, *cirbp*) and ribosomal genes, indicating the lack of a specific transcriptional programme (Fig. S3C, S4B). The basal cells across clusters (Bc1-4) expressed increasing levels of forkhead box TF (*foxa1*), ectoderm specifying TF (*tp63*), actin-binding proteins (*pfn1, tmsb4x*) and ion channel members (*ano1, kcna*) (Fig. 2C, S3C, S4C). The multiciliated cells across clusters (Mcc1-3) express increasing levels of structural (*tekt2*), cilia (*cfap45*), cytoskeletal components (*dynlrb2, tubb4b*) (Fig. 2C, S3C, S4D) and caveolin (*cav3*) that regulates cilia length and beat frequency (*2, 36*). The joint embedding and clustering identified an ionocyte and goblet cell cluster respectively. The osmoregulatory, proton-pumping ionocytes (Ic) expressed forkhead transcription factor I1 (*foxi1*), multimeric V-type ATPase family (*v1a, v1g, v0d*), cytochrome oxidase subunits (*cox7a2*) and carbonic anhydrases (*ca12*), catalysing protons and bicarbonate transfer (Fig. 2C, S3C, S4E), while the goblet cells (Gc) expressed angiopoietin (*angpt4*), lectins (*itln1*, *fucolectin*) and unconventional mucin glycoprotein otogelin (*otog,* also known as *mucXS* and *otogl2)* (Fig. 2C, S3C, S4F).

The high-dimensional integrated analysis captured all the major cell-types based on expected marker gene expression but also revealed novel insights. Firstly, the complete continuous MCE development could be integrated and visualised in low-dimensional space, capturing the cell state specialisation and transitions across branches. The homogeneous pluripotent cells undergo specialization at early neurula, forming highly heterogeneous early epithelial progenitor clusters (Fig. 2A-B). The broadly dispersed early epithelial progenitor states across the developmental manifold express stochastic levels of multiple markers, retain transcriptional plasticity and competence to respond to multi-lineage cues (by stages 18-20) and execute cell-type specific programs at tailbud stages. Secondly, we observed multiciliated progenitors by stage 13 (alongside early epithelial progenitors), marking an early priming event for ciliogenesis. Thirdly by stage 27, the basal cells developed alongside multiciliated, ionocyte and goblet cells, as the most abundant cell-type (*i.e.,* the proportion of cells/total sample proportion per stage; methods). Notably, the observed basal cell differentiation dynamics were different from other cell-types, owing to the presence of multiple late-stage subclusters and the expression of distinct TFs. We observed low levels of TF *tp63* within early epithelial progenitors (primed to form basal cells) and increased specifically across basal cell clusters at tailbud stages. These late-stage basal clusters were highly heterogeneous and constituted cells expressing either *tp63* alone or both *tp63* and *foxa1* (Fig. 2C). Lastly, standard community-based clustering approaches were suboptimal than graph-based methods (applied here) at deconvoluting the continuous MCE manifold, owing to irregular and highly continuous density of single-cells over MCE developmental time-course (Fig. S3D). Overall, the profiled scRNA-seq MCE developmental atlas and analysis reliably captured the developmental transitions starting from pluripotent stem cells to specialised mucociliary epithelial cell-types.

### Developmental trajectories and cell-type formation during MCE development

To characterise the developmental trajectories and maturation of different cell-types during MCE development, we *(i)* calculated independent pseudo-time ordering of all single-cells (Fig. 3A), *(ii)* computed single-cell differentiation potential (Fig. 3B), and *(iii)* inferred individual branch probabilities for pluripotent cells to reach different discrete cell-types (Fig. 3C).

**Fig. 3.**
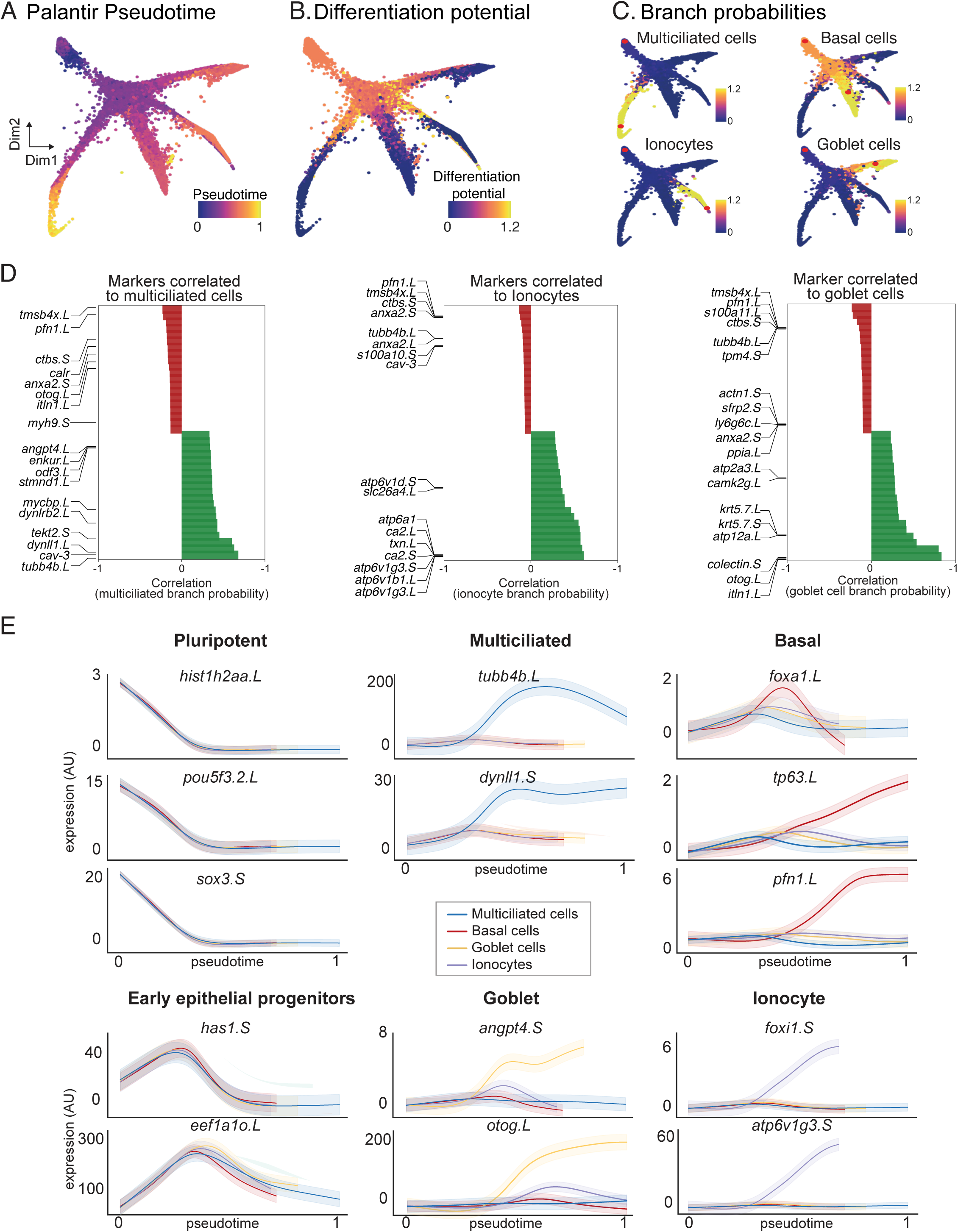
**Developmental transitions and cell-state branching over MCE trajectories** (A) Pseudotime inferred by Palantir overlaid on the MCE manifold (knn-graph). The pluripotent cells mark the beginning of pseudotime, while terminal cell states (Mcc, Gc, Ic and Bc) mark late pseudotime. (B) Differentiation potential (DP) overlaid on the MCE manifold (knn-graph) exclusively marks pluripotent and progenitor populations. (C) Branch probabilities assessed for each late-stage cell-type and overlaid over on the MCE manifold (knn-graph). The red dots mark the start (pluripotent) and respective end cells (cell-types). (D) Marker gene correlations to multiciliated, ionocytes and goblet cell branches of the MCE manifold (knn-graph). Green and red colours signify positively and negatively correlated genes respectively. (E) Gene expression trends of marker genes, using generalised additive models, contributing to individual branch probabilities including pluripotent (*hist1h2aa.L*, *pou5f3.2.L*, *sox3.S*), early epithelial progenitors (*has1.S*, *eef1ao.L*), multiciliated (pink: t*ubb4b.L*, *dynll1.S*), basal (green: *foxa1.L*, *tp63.L*, *pfn1.L*), goblet cells (orange: *angpt4.S*, *otog.L*) and ionocytes (blue: *foxi1.S*, *atp6v1g3.S*). The variable pseudotime reflects distinct cell-type diversification and differentiation speeds over MCE development.

We applied Palantir, to infer an unbiased pseudotime over the continuous high-density developmental manifold, as conventional approaches failed to generate cell-type specific trajectories. The pseudotime is inferred as a probabilistic process based on entropy over the entire manifold, with different maturation dynamics for each cell-type (Fig. 3A, E). As expected, we observed cells from blastula and neurula stages (pluripotent, early epithelial and ciliated progenitors) had low pseudotime, which increased across the branches consisting of specialised multiciliated, basal, ionocyte and goblet cells (Fig. 3A). The multiciliated branch had the highest pseudotime indicating that they were fully differentiated, relative to other cell-types (Fig. 3A).

As an independent measure of cellular plasticity, we calculated differentiation potential per single-cell (methods) and observed that pluripotent (P1, P2), ciliated (Mcc1) and early epithelial progenitors (Eep1- 4), had highest differentiation potential, drastically declining with lineage commitment across four branches (Fig. 3B). The differentiation potential strongly phenocopied the cell state assignment (Fig. 2A-B) and was inversely correlated with pseudotime (Fig. 3A-B), validating the cell states (subclusters) and cell-type assignment (multiciliated, basal, goblet, ionocytes) by multiple independent computational approaches.

As additional validation of cellular trajectories, we calculated branch probabilities of single-cells through iterative refinement of the shortest distance over the MCE manifold, assuming unidirectional progression and independence from neighbouring cells (Fig. 3C). The branch probabilities indicate the likelihoods of a pluripotent cell (red start cell at top) ending up as a differentiated and discrete cell-type (red end cell within multiciliated, basal, goblet and ionocytes, Fig. 3C, methods). The branch probabilities were low across pluripotent, early and multiciliated progenitors and dramatically increased across all late-stage cell-types, except basal cells (Fig. 3C). Remarkably, the branch probabilities of basal cells had a marginal increase (relative to multiciliated, goblet and ionocytes; Fig. 3C; top right), indicating that the basal cells share properties with pluripotent cells and act as late-stage stem cell reservoirs (*37, 38*). We further correlated the expression of known markers within individual branches and identified additional genes driving cell-type differentiation within single-cells (Fig. 3D). We observed that many genes had non-linear expression patterns and used generalised additive models to visualise averaged expression over respective cell-type pseudotime (Fig. 3E). The pluripotency markers (*pou5f3*, *sox3, hist1h2aa*) were highly expressed in early pseudotime across all cell-types and decreased with cell-type differentiation. Most cell-type-specific markers including multiciliated (*tubb4b*, *dynll1*), basal (*tp63*, *pfn1*), goblet (*angpt4*, *otog*) and ionocytes (*foxi1*, *atp6v1g3*), showed non-linear trends, selectively peaking at respective trajectories over the MCE pseudotime (Fig. 3E). Consistent with multipotent roles, the early epithelial progenitors peaked across all trajectories but sharply declined prior to specialisation (Fig. 3E). Notably, generalised additive models highlight that single-cell progression through the individual pseudotime lacked well-defined bifurcation points, further validating that early epithelial progenitors are plastic and undertake continuous cell-fate choices. Surprisingly, the expression dynamics of basal markers (*foxa1*, *pfn1*, *tp63)* strongly differed across pseudotime, highlighting heterogeneity and differential regulation of subclusters (Bc1-4) and spatial positioning within sensorial layers (Fig. 3E, 8B). The *foxa1* peaked in the middle (restricted to basal subcluster), while *tp63* and *pfn1* linearly increased over all basal cell pseudotime, distinct from other cell-types over MCE pseudotime (Fig. 3E). The generalised additive models confirm the robustness of the trajectory analysis (pseudotime, differentiation potential and branch probabilities) and further highlight that basal cell differentiation dynamics is differentiation to other MCE cell-types.

Next, we computed single-cell RNA velocities as a predictive model of differentiation dynamics, i.e., direction and speed of single-cell across complex developmental trajectories, by inferring a per-gene reaction model relating to abundance of unspliced and spliced mRNA (*39, 40*). The velocity estimates confirmed homogeneity in blastula, gastrula stages and progression through progenitors to the discrete cell-types at tailbud stages (*i.e.,* non-overlapping arrows within discrete clusters). The early epithelial progenitors across neurula stages (stages 13-20) were highly plastic (smaller arrows) and separated from multiciliated progenitors (Mcc1 to Mcc2 and Mcc3, Fig. S5A).

We next focussed on the characteristics of individual cell-types (Fig. S6-7). Zooming in on the early epithelial progenitors, we observed that single-cells within subclusters (Eep1-4) were not uniquely biased but had shared propensities towards ionocytes (*foxi1*, *ca2*, *bsg*), goblet (*otog*, *itln*, *fucolectin*) and basal cells (*tmsb4x, has1, ctbs*), confirming that early epithelial progenitors as a multipotent population (Fig. S6A-B). Contrary to early epithelial progenitors, the multiciliated cell differentiation followed stepwise progression starting from multiciliated progenitors (Mcc1, stages 13-20) and maturation into multiciliated cells (Mcc2 and Mcc3, tailbud stages, Fig. S6C-D). The tailbud-stage basal cell subclusters (Bc1-4, stages 22-27) located within the deep sensorial layers could be distinguished based on the expression of *tp63*, *pfn1*, *mal2* and *foxa1* (Fig. S7A-C). The Bc1 subcluster has the highest proliferating rates (S-phase) and is present throughout tailbud stages (similar proportion of cells, Fig. S7C). The proliferation rates decrease from Bc1 to Bc4 (decreasing S-phase, increasing G1-cells), indicating a reduced cell cycle speed. Both goblet and ionocytes underwent temporal specialisation from stage 22 to 24 (Fig. S7D-E). Intriguingly, we observed two separate ionocyte maturation trajectories across tailbud stages (Fig. S7E), which we further characterised as type I and type II ionocytes (Fig. 5D-E, S12A).

In summary, we capture the non-hierarchical mode of MCE lineage commitment that starts with the emergence of multipotent early epithelial progenitors that stochastically execute transcription programs within a defined developmental window (neurula stage), followed by maturation into different specialised cell-types (basal, ionocytes and goblet cells) at a similar developmental time, but with different dynamics and maturation speeds.

### Cell-type specific transcription factors and signalling pathways during MCE development

To identify the timing and order of transcriptional programs, we analysed expression dynamics of differentially enriched transcription factors (*41*) and signalling pathways across MCE development (methods, Table S6). The pluripotent clusters globally expressed the highest number of genes (Fig. 4A, D, E, S1C), involved in most cellular processes such as metabolic, cell cycle and repair (Gene Ontology (GO) terms, Fig. 4B-C). The early epithelial progenitors had few/no enriched genes, and shared GO terms with other cell-types. The multiciliated clusters were enriched with *cilia*, *microtubule-related* terms, while the ionocytes were enriched for ion transport and ATP hydrolysis (Fig. 4A-C). Most TFs were enriched across pluripotent clusters (pluripotent: *POU, SOX, yy1*, *ctcf, HMG* family; remodelers: SMAD, *arid2*, *jarid* family; activators: *znf326*, *znf706l*)), and downregulated with lineage specification (Fig. 4D, Table S6). We identified known and new cluster-specific TFs for multiciliated (*nsep1*, *ybx1*, *hes1*, *rfx2*, *id2*, *id4*, *sp7*), goblet cells (*grhl1*, *grhl3*, *nkx2-3*, *atf4*, *hic2*) and ionocytes (*akna*, *foxi1*, *dmrt2*, *tef*, *esrra*). The basal TFs included developmental (homeobox, caudal ParaHox, bHLH family), secretory (*foxa1*, *gfi1*, *tcf25*, *pbx2*) and growth regulators (*tp63*, *tbx3*, *tbx2*, *usf1*).

**Fig. 4.**
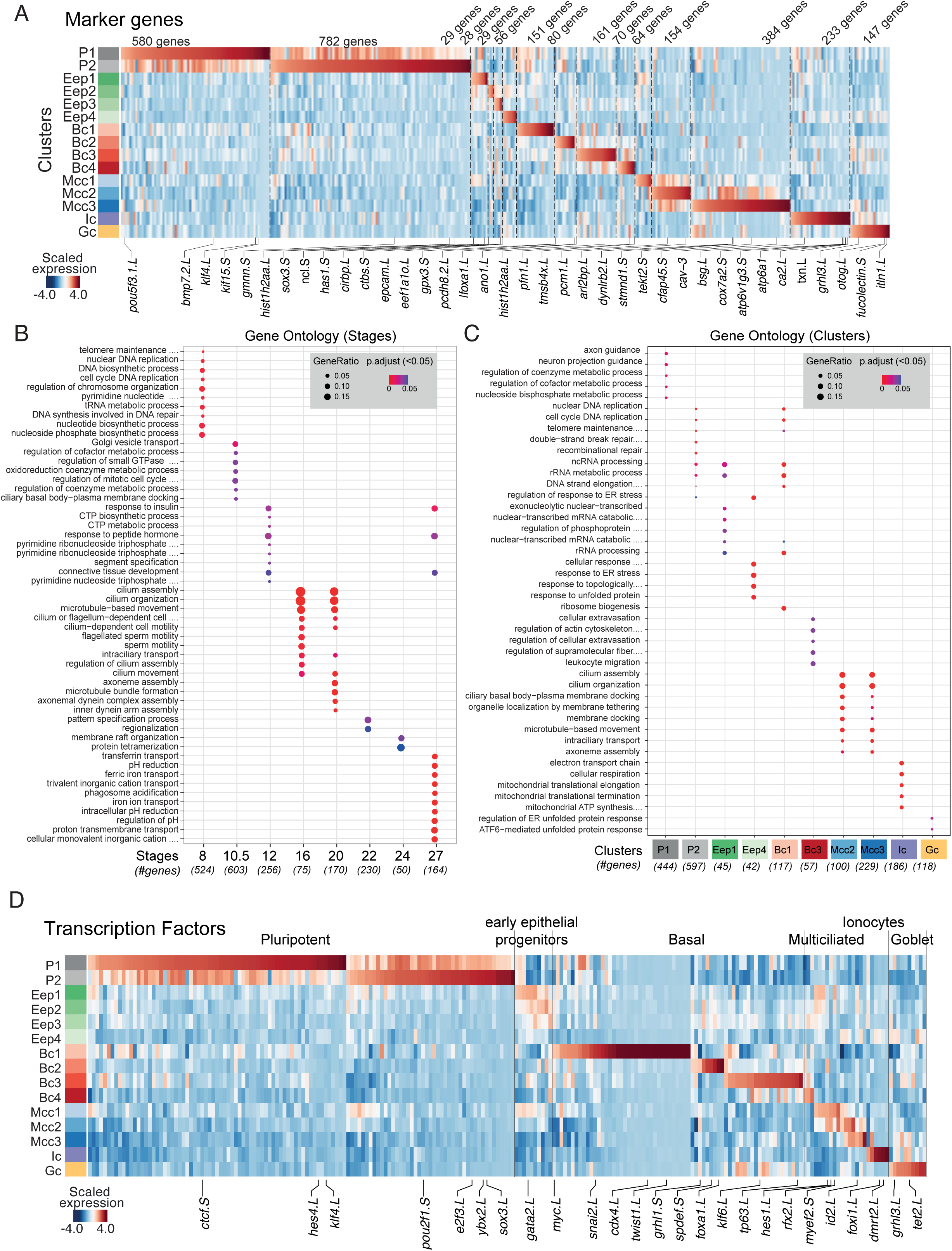
**Expression of marker genes, TFs and GO terms over MCE development** (A)) Heat map showing differentially expressed markers across MCE clusters. (B)) Significant Gene Ontology (GO; biological processes) terms for each MCE developmental stage. (C) Significant Gene Ontology (GO; biological processes) terms for each PhenoGraph cluster. Five clusters had no significant terms and were not presented. The gene ratio highlights the percentage of total GO term genes identified as enriched within each group (developmental stage/PhenoGraph cluster). (D) Heatmap showing the expression pattern of differentially expressed transcription factors (x-axis) over MCE PhenoGraph clusters (y-axis). The scale bars (A and D) indicate the z-scaled mean expression of respective markers.

Although blastula-stage organoids are cultured in absence of signalling factors and morphogens, several signalling pathways and gradients contribute to their default *ex vivo* specification into MCE cell-types. We observed enrichment of multiple non-canonical signalling pathways across pluripotent (Notch, Wnt and TGF-B, Fig. S8A, Table S7) and basal cell clusters, while the specialised cell-types (multiciliated, goblet and ionocytes) had fewer and highly restricted expression (Fig. S8A, Table S7). Notably, we observe stage-specific upregulation of Notch ligands and receptors in basal cell subclusters (*notch2, smad3, smad7*) and ciliated progenitors (Mcc1: *hes1, hes8*) (Fig. S8A, Table S7). We next compared lineage-specific TFs and known marker genes identified from bulk transcriptomics (*42*), and observed a striking single-cell heterogeneity, particularly across multiciliated cells, ionocytes and basal cells (Fig. S8B-D, Table S6-S7, supplementary note 2). The reported multiciliated markers at single-cell level had a stepwise activation from Mcc1 (progenitors) to Mcc3 (mature), averaged and missed across bulk assays (Fig. S8B), while only a subset of bulk ionocyte markers was strongly enriched at single-cell level across ionocytes (Fig. S8C) (*21*). We observed a similar pattern with recently published basal cell markers (Fig. S8D) (*21*). The expression heterogeneity at single-cell level and discordance with bulk expression indicate that TF activity within subclusters is tightly coupled with cell-type maturation, and cell-type specific TFs have a high degree of positive regulation during MCE development.

Intrigued by the expression discordance of bulk markers at single-cell level, we investigated how molecular features (expression noise, coefficient of variation, transcript diversity) varied over MCE development. Within populations, individual cells have been reported to resolve cellular decisions between stemness and uncertainty through increased heterogeneity and changes across molecular features to drive noise-induced differentiation (37, 38). We observed that both the coefficient of variation and entropy increased over MCE development and was highest at the tailbud stages (Fig. S9A-B). Notably, transcriptional diversity (number of unique transcripts) was anti-correlated and decreased as single-cells underwent lineage commitment (Fig. S9A-B), indicating an inherent stochastic gene expression within neurula stage progenitors contributing to their plasticity through the expression of multi-lineage expression programs (low expression of many unique transcripts). The progression to late stages leads to the execution of specific cell-type gene regulatory networks (increased expression of fewer specific transcripts; higher CV, low transcriptional diversity) (*43*). We also re-analyzed the MCE atlas through the recent CytoTrace framework, which measures expressed genes per cell as a proxy for transcriptional diversity and cell-type differentiation (*44*). We confirmed that early and multiciliated progenitor clusters (Eep1-4, Mcc1-2) had a low CytoTrace score indicating multipotent characteristics, while the late-stage cell-types (Bc1-4, Mcc3, Ic and Gc) had the highest scores indicating a mature state (Fig. S9C-D), further validating that MCE developmental decisions are modulated through changes in molecular features. The summarised comparison of different molecular features is highlighted in Fig. S9E (Tables S1, S9). Taken together, our analysis suggests that the neurula stages provide the developmental window for individual cells to exhibit different cell fate-switching frequencies and establish initial heterogeneities that are resolved by cell-type specific regulatory networks.

### *In silico* cell lineage inference and in-situ validation of cell-type dynamics across MCE development

Given our time-resolved MCE developmental scRNA-seq atlas, we devised an *in silico* lineage inference algorithm to map cell state and cell-type relationships (Fig. 5A-D); validated their temporal emergence and dynamics in whole embryos using *in situ* hybridization chain reaction (HCR) and multiplex spatial imaging. While lineage inference from scRNA-seq data is non-trivial, we devised a generalizable method built on a neighbourhood mapping and consensus voting strategy, requiring few/no priors to map the single-cells and their cluster relationships between successive developmental stages, and charted the entire MCE development (Fig. 5A-B, methods, supplementary note 3). The inferred lineage tree provided several insights into developmental timing, TFs, marker gene expression, cell cycle and state relationships over MCE development (Fig. 5A-C).

**Fig. 5.**
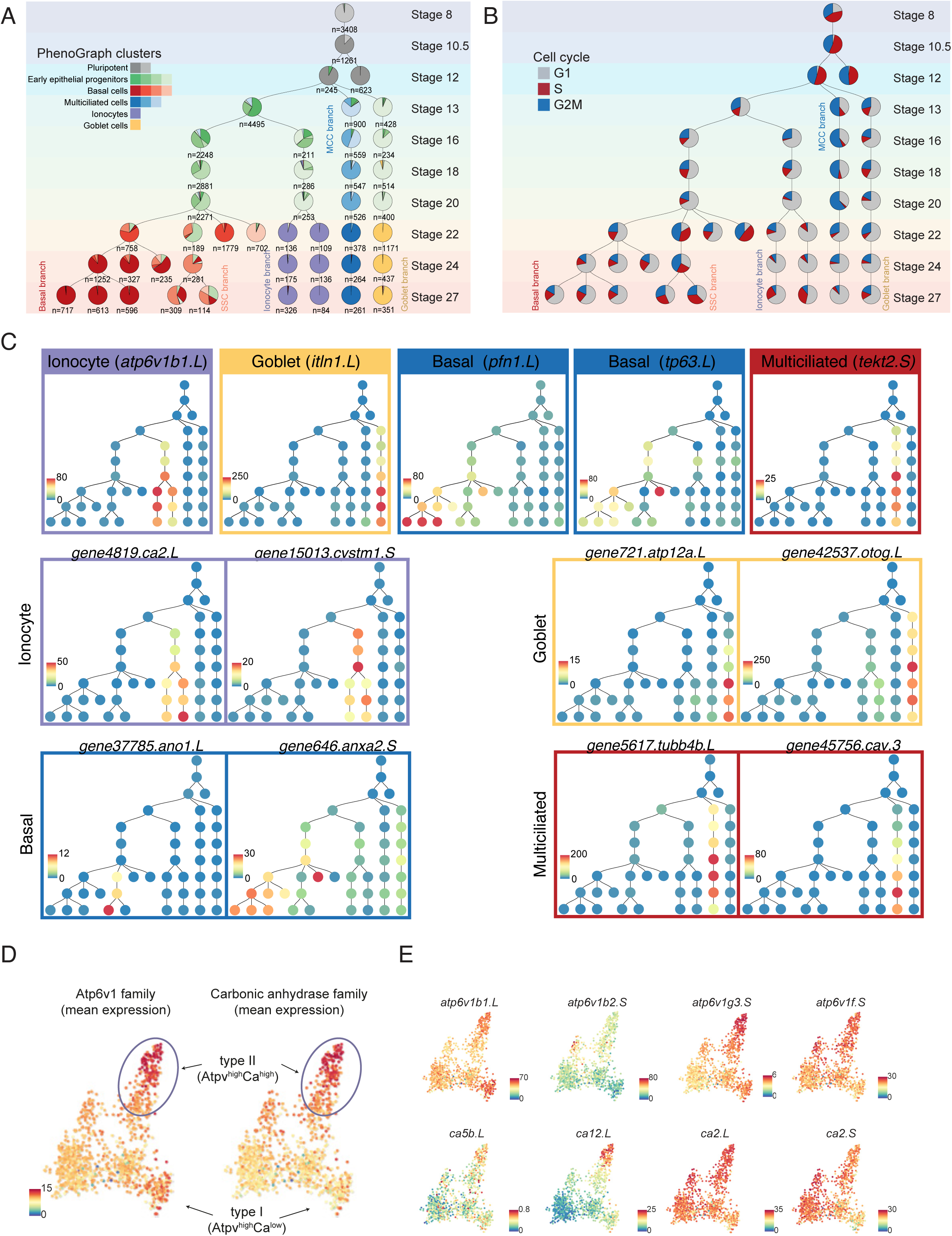
**Lineage inference of developing MCE cell-types.** (A) *in silico* lineage inference of the cell states and types across developing MCE. The number of cells within each group are highlighted underneath each node. The pie chart indicates proportions of cells mapping to clusters (consensus voting, see methods) (B) Lineage inference map overlaid with proportions of cells across cell cycle stages. (C) Timing and expression of different marker genes (Ionocytes: *atp6v1b1.L*, Goblet: *itln1.L*, Basal: *pfn1.L, tp63.L,* Multiciliated cells: *tekt2.S*) over the inferred lineage tree. Lineage restricted expression of additional markers of ionocytes (*ca2.L*, *cystm1.S*), goblet (*atp12a.L*, *otog.L/mucXS/otogl2.L*), basal (*ano1.L*, *anxa2.S*) and multiciliated cells (*tubb4b.L*, *cav3.S*). (D) Low dimensional visualisation of ionocyte subpopulations, overlaid with the mean expression of Atp6v1 and carbonic anhydrase family members, indicating type-I (Atp6v1^high^Ca^low^) and type II ionocytes (Atp6v1^high^Ca^high^). (E) Low dimensional visualisation of ionocyte subpopulations overlaid with expression of few Atp6v1 and carbonic anhydrase family members. The scale bars indicate the scaled imputed expression of respective markers.

Firstly, the multiciliated transcriptional programme was initiated directly from pluripotent tissue as a distinct lineage (*mcidas, tekt2* expression at stage 13) and underwent maturation into multiciliated cells, with decreasing cell cycle speeds and proportionally fewer cells, relative to the total cellular pool (blue branch, Fig. 5A-C). The single cell *in situ* HCR in embryos confirmed the timing and lineage activation of multiciliated progenitors (*tekt2)* by stage 13, mutually exclusive with other cell-types (Fig. 6A-C, S10A). We observed a continuous increase of the *tekt2* mRNA fluorescent signal in the cytoplasm as multiciliated progenitors radially intercalated (*i.e.,* apical movement) from the deep sensorial layers and integrated within the superficial epithelium at stage 24 (Fig. 6A-C, S10A).

**Fig. 6.**
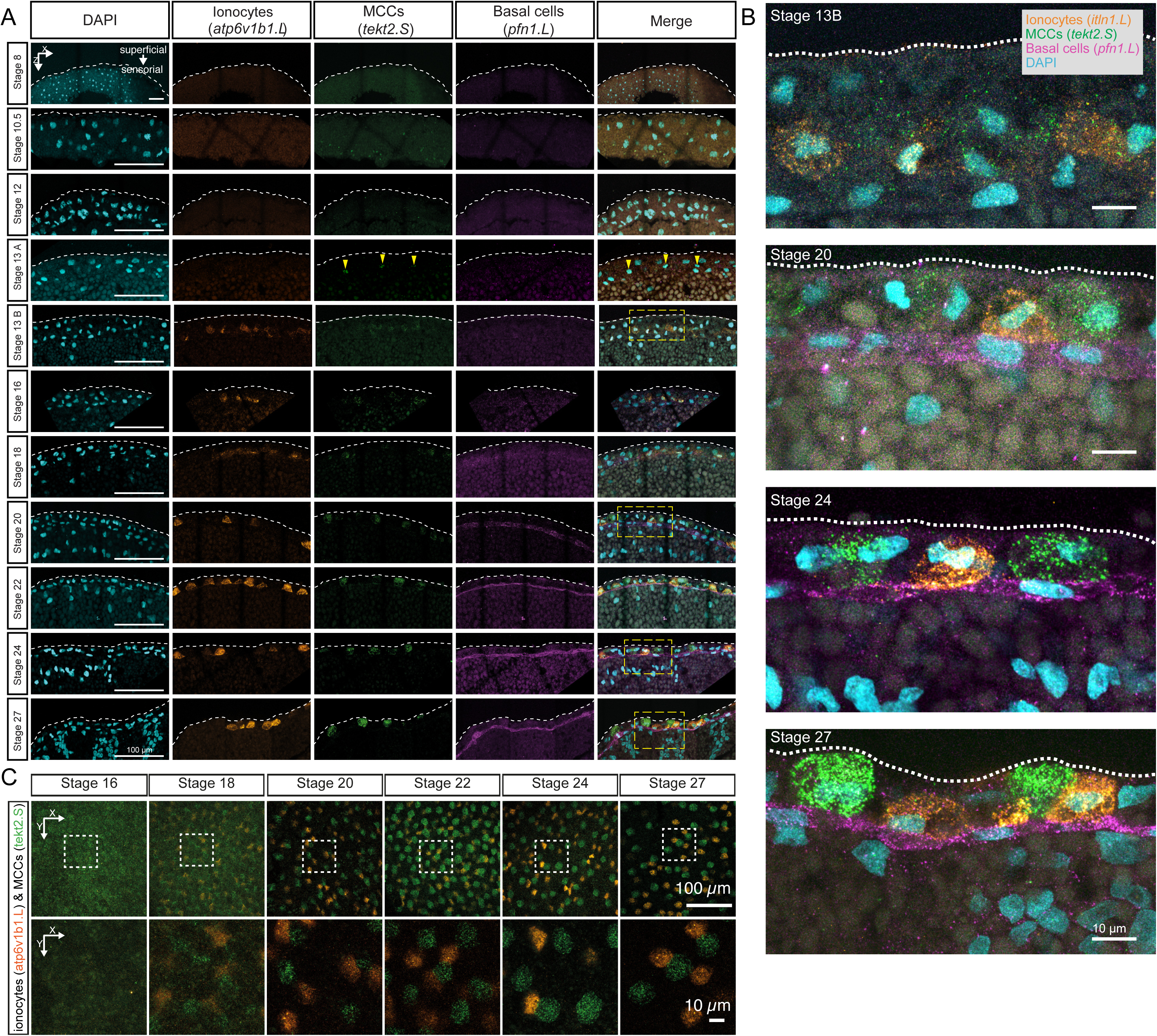
***In-situ* HCR validation of multiple cell-types lineages over MCE development** (A) *In-situ* hybridization chain reaction (HCR) and validation of lineage inference over 10 development stages of the embryonic epidermis in embryos, marking ionocytes (*atp6v1b1.L*, orange), multiciliated (*tekt2.S*, green) and basal cells (*pfn1.L*, magenta). The nuclei are marked by DAPI staining (cyan). The multiciliated cells and ionocytes emerge during neurula in the sensorial layer and radially intercalate (move apically) to superficial layers by early tailbud stages. Multiciliated cells are consistently observed at stage 13 (panel - stage 13A), whereas ionocytes are consistently observed at stage 16; in four out of nine embryos, ionocytes are observed at stage 13 (panel - stage 13 B). The basal cells are retained in the sensorial layer. The yellow rectangles indicate zoomed-in regions shown in panel B. Yellow arrowheads point to regions (within the nucleus) expressing *tekt2.S*. (B) Zoomed view of i*n-situ* HCR marking expression, positioning and migration of multiciliated (*tekt2.S*, green), ionocytes (*atp6v1b1.L,* orange) and basal cells (*pfn1.L*, magenta). Images A and B represent maximum intensity projections of transverse cross-sections and white dotted lines indicate apical superficial epithelium. (C) *In-situ* HCR of whole embryonic surface epithelium marked by multiciliated (*tekt2.S*, green) and ionocytes (*atp6v1b1.L,* orange) at the superficial epidermis. Multiciliated cells and ionocytes move apically (radially intercalate) from the sensorial to the superficial epithelium and distribute in a salt-and-pepper fashion. Images represent maximum intensity projections of Z-sections taken from the apical superficial epithelium. The white rectangles indicate zoomed-in regions shown in panel C, lower row.

Secondly, the secretory transcriptional programme was established exclusively through the early epithelial progenitors and their maturation, in contrast to the proposed early-stage basal cells in human and mouse cells (*10, 45*). The goblet-primed early epithelial progenitor lineage could be observed as early as stage 13, with a sporadic expression of marker genes (*otog/mucXS/otogl2, itln1*, Fig. 5A-C), but underwent maturation only at tailbud stages (yellow branch, light green pie, Fig. 5A-C). Distinct from goblet cells, the ionocytes differentiated from a separate early epithelial progenitor pool by stage 16, and also underwent maturation at tailbud stages (purple branch, mixed green pie, Fig. 5A-C). Using *in situ* HCR, we validated the consistent expression of ionocytes markers at stage 16 embryos (*atp6v1b1*: cytoplasmic, Fig. 6A-B, S10A). Notably, in a few embryos (4 out of 9), we observed a few cells (presumptive Eep) that expressed ionocyte markers within single-cells already at stage 13, which we confirmed both by scRNA-seq and *in situ* HCR (*atp6v1b1;* Fig. 6A-B, S10A). The presence of these cells at stage 13 further indicates that neurula stages provide the temporal developmental window for single-cells to stochastically drive multi-lineage programs, which however, are executed at tailbud stages. Similarly to *tekt2* in multiciliated cell progenitors, *atp6v1b1* cytoplasmic expression increased within superficial epithelium as development progressed, efficiently labelling radially intercalating ionocytes distributed in salt-and-pepper fashion next to the multiciliated cells (Fig 6C). The developmental dynamics and transcription factors that drive goblet cell differentiation remain largely unresolved (Fig. 7A). Using *in situ* HCR, we validated the faint, interspersed expression of goblet markers in stage 13 embryos increasing over the development (*otog/mucXS/otogl2, itln1*, Fig. 7B-D, S11A, whole-mount embryos) marking both the nucleus and the cell outline. The non-uniform distribution of *otogl2* and *itln1* was particularly prominent within the apical surface of the superficial epithelium, outside the neuroepithelial plate (not marked by *otogl2* nor *itln1*), at stage 13 transitioning to more homogeneous expression from stage 20 onward (Fig. 7C, S11), which parallels the reported maturation of goblet cells upon regeneration (*46*).

**Fig. 7.**
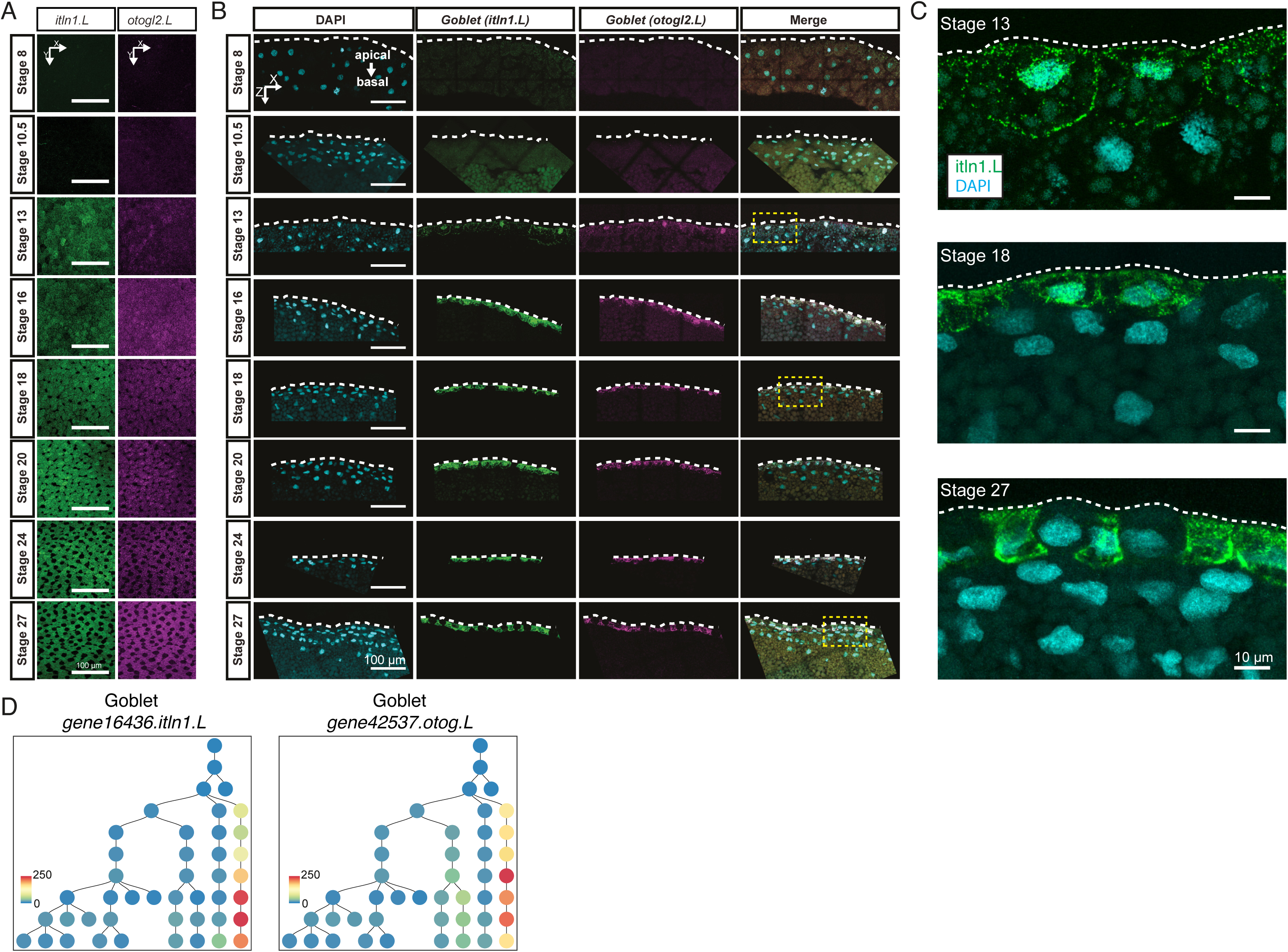
***In-situ* HCR validation of developing goblet cell lineage over MCE development** (A) *In-situ* HCR of whole apical superficial epithelium indicating maturation of goblet cells (*itln1.L*, green) and otogelin (*otog.L/mucXS/otogl2.L*, magenta). Dark cells (black hollow spaces) from stage 18 (here, panel C) represent radially intercalating multiciliated cells and ionocytes specified in the deep epithelial layer. Images represent maximum intensity projections of Z-sections taken from the apical superficial epithelium. (B) *In-situ* hybridization chain reaction (HCR) of 8 developmental stages indicating goblet cell differentiation marked by intelectin (*itln1.L*, green) and otogelin (*otog.L/mucXS/otogl2.L*, magenta); nuclei are marked by DAPI staining (cyan). The yellow rectangles indicate zoomed-in regions shown in panel C. Images (here and in C) represent maximum intensity projections of transverse cross-sections and white dotted lines indicate apical superficial epithelium. (C) Zoomed-in view of *in-situ* HCR marked by goblet cells (*itln1.L*, green) and otogelin (*otog.L/mucXS/otogl2.L*, magenta). At stage 13, the goblet markers are detected at very low levels with interspersed staining; but gradually mark both nuclear and cytoplasm by stage 22. (D) Timing and expression of goblet marker genes (*itln1.L*, *otog.L*) over the inferred lineage tree. The scale bars indicate the scaled imputed expression of respective markers.

Thirdly investigating ionocyte lineage, we classified *foxi1-marked* ionocytes into type-I (*Atp6v1^high^Ca^low^*) and type-II (*Atp6v1^high^Ca^high^*), distributed in a salt and pepper pattern across embryonic MCE (Fig. 5D- E, 6A-C, S12A) (*47, 48*). The ionocytes homeostasis and regeneration are regulated by *cftr*, and its mutations cause osmotic imbalance and modified mucus properties leading to cystic fibrosis (*10, 23*). While *cftr* expression was missing in our single-cell dataset, we reconfirmed its patched expression in *foxi1+* ionocytes using bulk transcriptomics data and *in situ* hybridization at tailbud stages (Fig. S12B- F) (*49*).

Fourthly, the actively cycling basal cells (*tp63, pfn1*) formed multiple subclusters (Fig. 5A-C). The mammalian basal cells act as adult stem cells replenishing airway epithelial cells, but their function in *Xenopus* MCE has remained elusive (*9, 50*). The *in situ* HCR validated basal cells (*pfn1.L, tp63.L)* appearance by late neurula, progressively concentrating within the sensorial layers during development (Fig. 6A-B, 8A-B). The abundance of cycling basal cell subpopulations strongly suggests that the turnover of exhausted specialised cells occurs through the differentiation of late-stage basal clusters (*51, 52*). Notably, we observe expression of *foxa1* by stage 22, labelling flat cells sitting on top of the basal cell layer expressing *tp63* (Fig. 8B). *Foxa1* is thought to mark the serotonin secretory cells at the tadpole stage (stage 32), and intriguingly, we did not observe other secretory markers. During later stages, the morphology of *foxa1* expressing cells becomes rounded alongside migration toward the superficial layer (Fig. 8B), consistent with their radial intercalation into the *Xenopus* epidermis (*53, 54*).

**Fig. 8.**
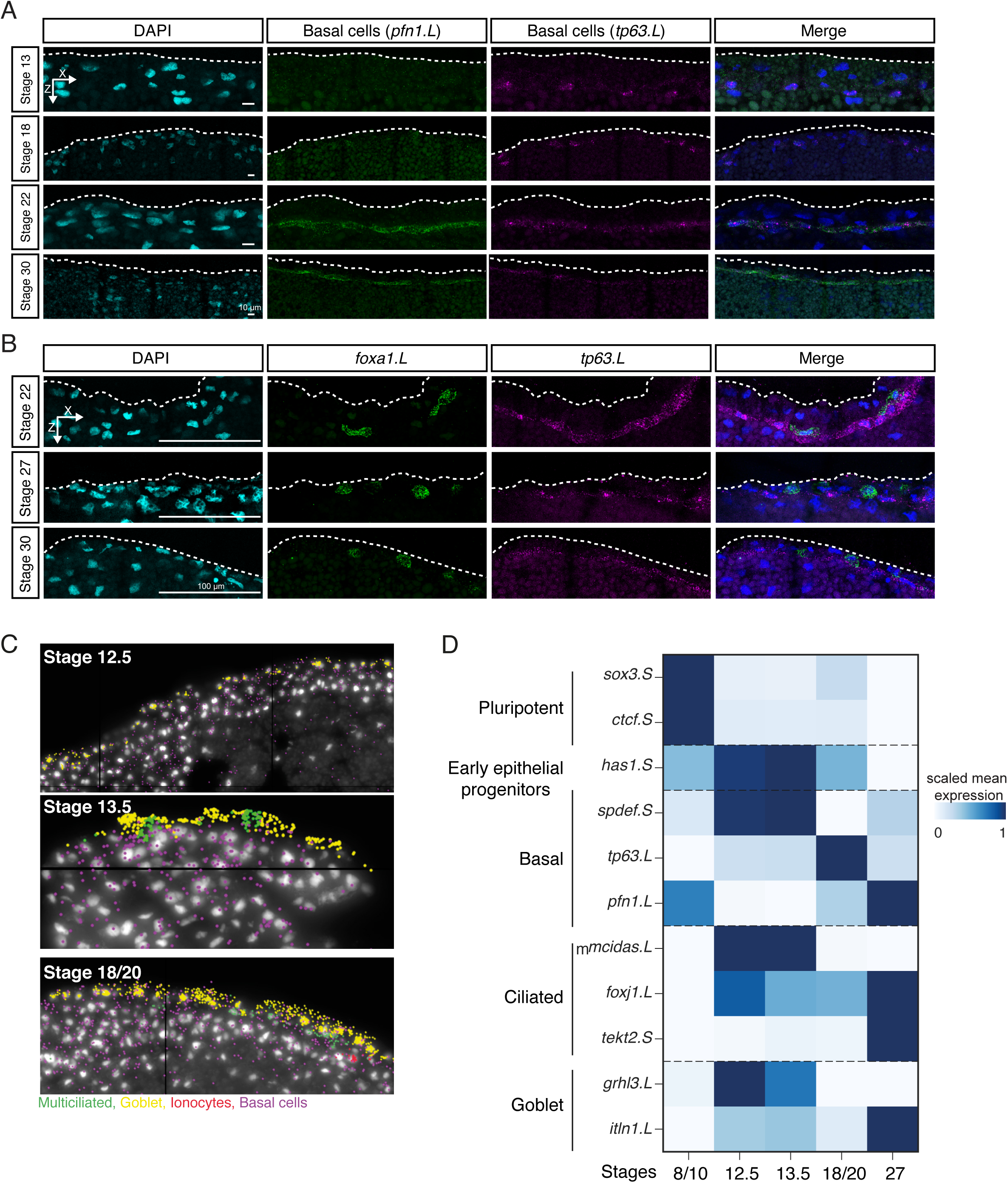
***In-situ* HCR validation of developing basal cell lineage over MCE development** (A) *In-situ* hybridization chain reaction (HCR) of apical superficial epithelium indicating basal cells marked by Profilin (*pfn1.L*, green), Tumour protein P63 (*tp63.L*, magenta) across neurula (stage 13 and 18) and tailbud stages (stages 22 and 30); nuclei are marked by DAPI staining (in cyan, separate channel and in blue, merged channels). The expression levels of *pfn1.L* and *tp63.L* decrease by stage 30. Images (here and below) represent maximum intensity projections of transverse cross-sections and white dotted lines indicate apical superficial epithelium. (B) *In-situ* hybridization chain reaction (HCR) marked by forkhead box A1 (*foxa1.L*, green) and tumour protein P63 (*tp63.L*, magenta) across tailbud stages. (C) Representative images of single-cell multiplexed RNA imaging in whole embryos across 3 MCE developmental stages. Stages 12.5 and 13.5 mark the transition into neurula development. (D) Heatmap showing the RNA abundances of different cell-types markers, as captured by multiplexed single-cell RNA imaging. The early epithelial progenitors are marked by the expression of multiple markers at early stages, while specialised cell-types exclusively express markers at late stages. The scale bars indicate z-scaled mean expression in aggregated cells.

Lastly, through multiplexed single-cell RNA imaging in whole embryos, we re-confirmed the transition into gastrula and neurula stages (stages 12.5-13.5) is accompanied by loss of undifferentiated state marker expression (*sox3.S*) and the presence of early epithelial progenitors (*has1.S, spdef.S, grhl3.L*) and multiciliated progenitors (*mcidas.L, foxj1.L*) (Fig. 8C-D). The radially intercalating multiciliated (*foxj1.L, tekt2.S*) and goblet cells (*grhl3.L, itln1.L*) underwent respective maturation, while the basal cells within sensorial layers were marked by heterogeneous marker gene expression (*tp63.L, pfn1.L*) (Fig. 8C- D).

Taken together, the lineage inference, *in situ* HCR and multiplex RNA imaging at single-cell level provides a detailed understanding of how cell-fate decisions (i.e., early epithelial progenitors) are undertaken within a temporal developmental window (i.e., neurula stages). We provide evidence that cell-fate decisions, including the emergence of cell states and their underlying cellular plasticity during MCE development, are driven by a combination of molecular features (expression variance, entropy, noise) (*55, 56*) and deterministic qualitative changes (changes in cell shape and migration accompanying radial intercalation) (*57, 58*).

### Identification of early epithelial progenitors across *Xenopus* species

To validate that early epithelial progenitors are a bonafide cell-type that drive secretory (and basal cell) lineage programs, we mined the single-cell *Xenopus tropicalis* developmental atlas (*24*), comparing the matched non-neural ectoderm lineage (stages 8-22) with our *Xenopus laevis* MCE developmental atlas (stages 8-27) (Fig. 9A-D, S13A-F). While the late-stage cell-types were shared between the atlases (*i.e.,* Stage 22), the lack of early epithelial progenitors at neurula stages was puzzling. We initially correlated known cell-type markers over MCE development and observed that early epithelial progenitor subclusters expressed secretory and basal markers at lower levels than late stage cell-types (Fig. S14). Notably, the early epithelial progenitor subcluster (Eep4) expressed both goblet and ionocyte markers. Therefore, we classified a conservative secretory signature (33 goblet and 20 ionocytes markers) that robustly marked both cell-types across both atlases (Fig. S13A, D). Using the signature gene sets, we confirmed that both mean expression and number of cells of goblet cells and ionocytes, respectively, increased over development across both studies (Fig. S13B, E). Surprisingly, many cells within the *X. tropicalis* atlas expressed both secretory signatures (Fig. S13C, F), as observed with early epithelial progenitor subclusters (Fig. S14). We indeed confirmed that these double-positive cells (Ic^+^, Gc^+^) were enriched within early neurula stages and visualised single-cells in a low dimensional expression space (Fig. 9A). Notably, the double-positive cells (Ic^+^, Gc^+^) expressed higher levels of goblet markers (than ionocytes) and were likely classified as goblet cells. We further confirmed that early epithelial progenitors (this study) are indeed correlated with the double positive cells (Ic^+^, Gc^+^) at neurula stages (*X. tropicalis* atlas) (Fig. 9C, D), while the late-stage secretory cells (this study) correlate with late-stage ionocyte and goblet cells (*X. tropicalis* atlas) (Fig. 9C, D). We performed *in situ* HCR with ionocyte (*atp6v1b1*) and goblet marker (*otogl2*) in neurula stage embryos, and found double-positive cells in both sensorial and superficial layers. The sensorial cells expressed higher levels of *atp6v1b1*, while superficial cells had higher *otogl2*, respectively, indicating distinct subpopulations (Fig. S13G). In summary, MCE developmental progression requires transition through the bonafide multipotent early epithelial progenitors prior to differentiation into the secretory cell-types, and the early epithelial progenitors have likely been misannotated owing to variable expression of multiple secretory markers.

**Fig. 9.**
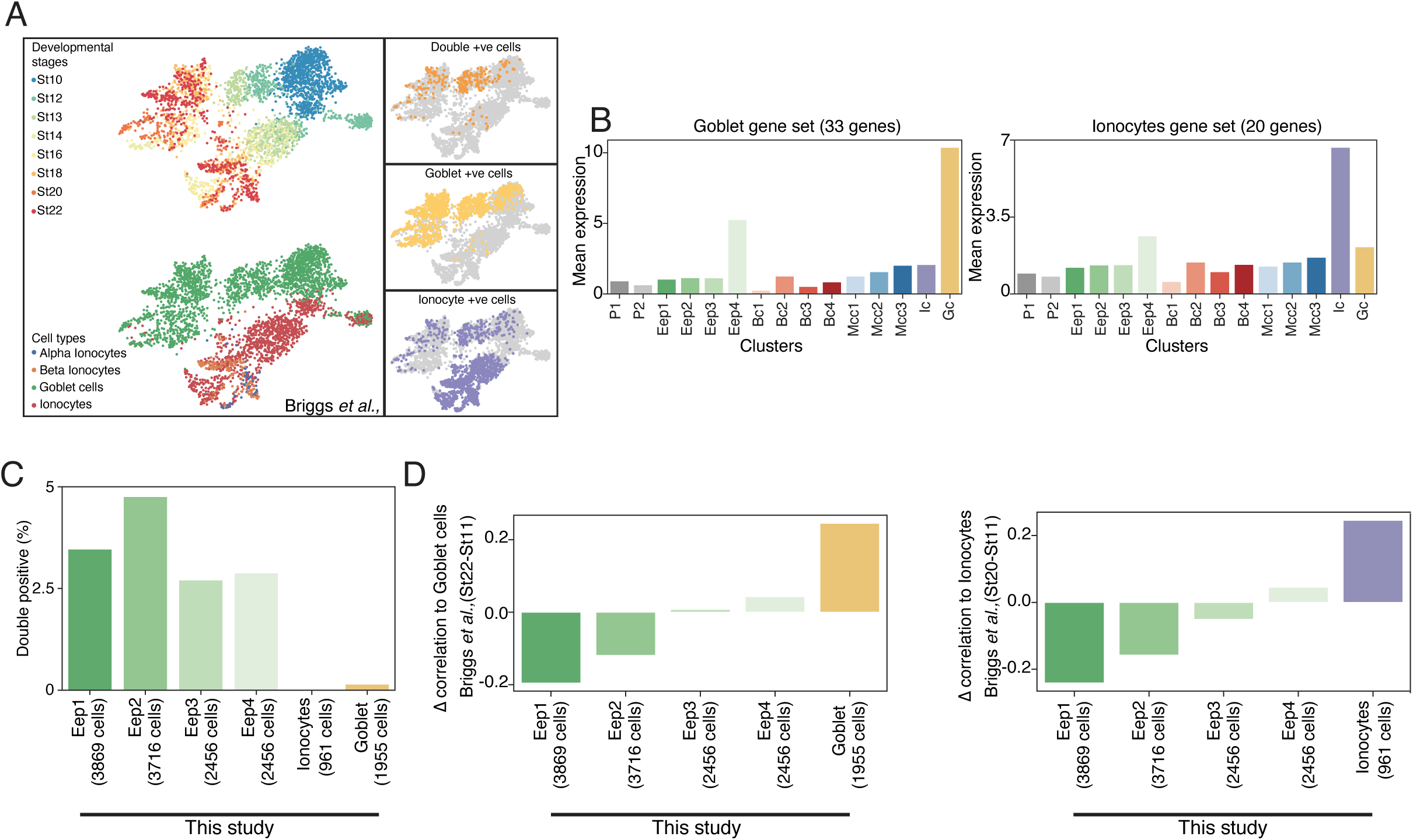
**Function of multipotent early epithelial progenitors in the development of the secretory cell-types.** (A) Low dimensional visualisation of non-neural ectoderm cells, stages and their annotations across *X. tropicalis* development atlas. The insets indicate the goblet cells, ionocytes and double-positive cells marked by core signature gene sets. (B) Mean expression of a goblet (33 genes) and ionocyte (20 genes) signature genes over MCE developmental clusters (this study). While late-stage goblet cells and ionocyte have the highest mean expression of signature gene sets, the early epithelial progenitor clusters also express both signature genes. (C) Percentage of early epithelial progenitors expressing both goblet (33 genes) and ionocyte (20 genes) signature genes. Nearly all double-positive cells are early epithelial progenitors. (D) Change in the correlation between stage 22 and stage 11 goblet cells and ionocytes across *X. tropicalis* development atlas, and this study. The correlation at stage 11 (early stages) is higher to early epithelial progenitors, while at late stage (stage 22) correlation indicates respective goblet cells and ionocytes.

### Evolutionary conservation of MCE cell-types

Recently, many cell-types of the *Xenopus* MCE and their functional equivalents have been identified during airway development in mouse and humans, aided by single-cell transcriptomics (*10, 11, 20, 23, 59*). We investigated whether the transcriptional programmes (i.e., expression modules) were conserved across vertebrates and drive cell-type function. To characterise the conservation and divergence of MCE cell-types, we performed a comparative analysis of our data with eight single-cell transcriptomics atlases spanning *Xenopus laevis* (adult epidermal regeneration), *Xenopus tropicalis* (embryonic, adult epithelium), mouse trachea and human nasal epithelium (10, 23, 24, 45, 60–62), spanning 144 cell-types across 120,842 single-cells. To compare cell-types across different species, we re-analyzed all single-cell atlases in a standardised and uniform manner, retaining respective author annotations (grey shapes, Fig. 10A). We devised a differential expression-based cell-type enrichment score (Enrichment score, Fig. 10A, methods) for each cell-type (accounting for the number of cells; black rectangular bars DE genes, Fig 10A, methods) that robustly distinguished the secretory, ciliated and basal transcriptional cell-types from other cell-types, using author specified cell-types as ground truth (Fig 10A, S15A-D, supplementary note 4, methods). The ‘ciliated’, ‘secretory’ and ‘basal’ enrichment scores (red squares, Fig 10A) were calculated for each cell-type (across 9 single-cell atlases), indicating the specificity of marker genes to classify respective cell-types (Fig. S15A), enabling a comparative analysis between species. We observed the highest differentially expressed genes (DE; and respective scores) in ciliated cell-types, followed by basal and secretory cell-types (enrichment scores in red squares; the number of DE genes in grey bars Fig. 10A). The secretory cell-types (goblet cells, ionocytes and subtypes) from 3 *Xenopus* atlases were grouped together, but separated from their mouse and human counterparts (Tuft, Brush, PNEC, etc., cell-types) (grey shapes Fig. 10A, S15B). The *Xenopus* goblet cells lack canonical mucins present in higher vertebrates, while ionocytes were originally characterised in *Xenopus* (*47*) and their disease-related counterparts have been identified in the mammalian airway (*63, 64*). Similar to the secretory programme, the mouse and human specialised basal cell-types were grouped separately from both *Xenopus* counterparts (Fig. 10A, S15C), indicating a distinct higher vertebrate basal expression programme. Notably across ciliated cells, we observed a conserved transcriptional programme of ciliated progenitors across species (two mouse ciliated progenitors, three *Xenopus* ciliated progenitors) and clear distinction of mature ciliated cell-types (multiple mouse, human and *Xenopus* cell-types, Fig. 10A, S15D), indicating that paradigms of ciliogenesis and maturation are conserved across higher vertebrates. We highlight enriched genes and their expression in different cell types across species (Fig. S15E). In summary, the comparative analysis captured the functional importance and conserved transcriptional programmes during ciliogenesis across vertebrates, while the regulation of cellular identity across secretory and basal cell programs is driven by cell-type specific programs.

**Fig. 10.**
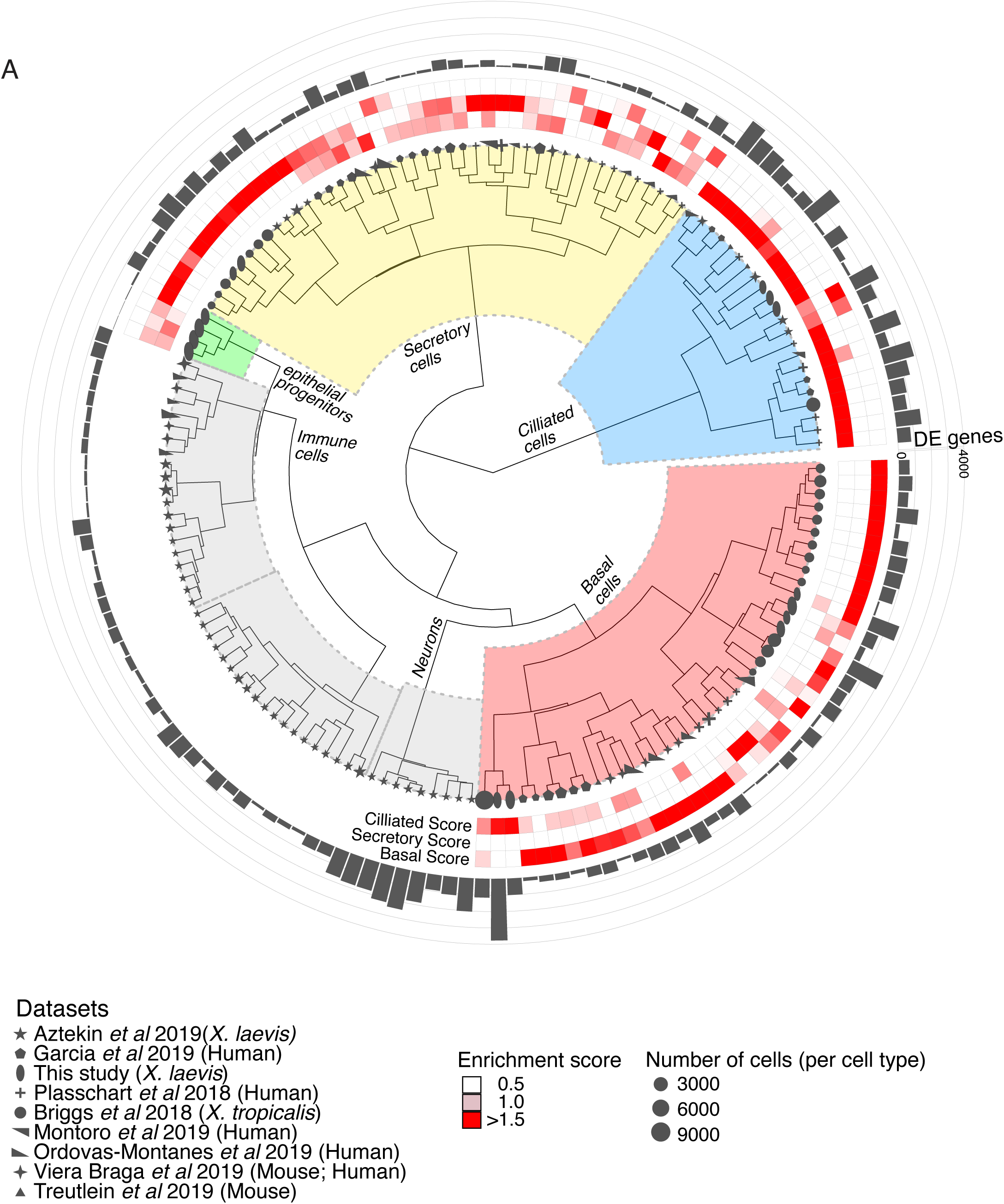
Comparative analysis of MCE cell-types and development. **(A)** Comparative analysis between *Xenopus*, mouse and human mucociliary cells from 9 single-cell airway atlases, using expressed orthologous gene sets. Using author-annotated cell-type labels (shapes), a cell-type enrichment score (ciliated, secretory and basal, enriched scores in red) is calculated based on differentially enriched genes (grey bars), which classified the different cell-types into Basal, Ciliated, Secretory and other cell-types (Immune, neuronal). The four early epithelial progenitor clusters together with secretory cell-types. The shapes indicate the respective study, shape sizes indicate the scaled number of cells within each author annotated cell-type, grey histograms indicate the number of differentially enriched genes per author annotated cell-type. The legend further indicates the respective study, organism type and species; colouring of enrichment score and sizes indicates the number of cells per author annotated cell-type.

## DISCUSSIONS

Investigating the composition, development ordering and dynamics of MCE cell-types during embryogenesis is of fundamental importance to vertebrate biology. The *Xenopus* mucociliary epidermis enables an understanding of mucociliary development without the prerequisite of lung formation and provides a critical developmental and evolutionary perspective to comprehensive studies in mammals focusing on MCE regeneration and homeostasis.

Here, we profiled functional MCE formation across ten developmental stages at single-cell resolution and captured transitory and mature cell-types over the MCE development that lack discrete bifurcation points. Our data highlights that homogeneous blastula undergoes the key initial developmental transition to the multiciliated progenitors and a multipotent early epithelial progenitor ensemble that drives the body plan (gastrula, neurula) and subsequent organogenesis in trajectories to specialised cell-types during tailbud stages. We report that *Xenopus* MCE differentiation follows a continuous non-hierarchical developmental model, which is neither discrete nor completely stochastic. We speculate that the two gastrula subpopulations represent the two-layered epidermal cells with varied plasticity in organoids and embryos. Furthermore, in this study, we uncovered the initial fate transition, alongside expression changes in TFs and marker genes, involved in the complex coordination of signalling, cell cycle and spatiotemporal cell-type organisation. The expression dynamics over the branched developmental manifold and pseudotime highlight that diverging multi-lineage programs occur at different developmental speeds. This dynamic equilibrium of progenitor specialisation is critical for maintaining signalling and maternal gradients, as the depletion of either multiciliated, ionocytes, basal or goblet cells leads to failures in the formation of terminal cell-type and mucociliary functions (*30, 65*). Through radial intercalation, multiciliated precursors migrate from the sensorial layer to embed between secretory cells at superficial epithelial layers (*66*). Alongside the maturation of multiciliated cells, we capture and validate early epithelial progenitors as the precursors for basal and secretory cellular programs. We observed heterogeneity in basal cell clusters, based on marker genes but also cell cycle proportions, which can contribute to regenerating adult MCE cell-types, as in mammals (*9, 50*). We uncover basal cell subpopulations, which retain the developmental plasticity to further differentiate and potentially replenish adult MCE cell-types. The *Xenopus* mature cell-types have recently been reported to undergo transdifferentiation to other cell-types (*67*). We speculate that this fate conversion could be facilitated through the early epithelial progenitors and subsequent basal cell subpopulations that retain cellular plasticity and maintain MCE composition in homeostasis and regeneration (*60, 67*).

Through MCE development, we observe stochastic gene expression within neurula-stage progenitors contributing to the expression of multi-lineage expression programs, resolving into late-stage cell-type specific gene regulatory networks. The molecular features (decreasing transcriptional diversity, entropy and increased molecular noise) coupled with cell shape and migration, indicate that neurula stages provide the developmental window to establish initial heterogeneity and cell fate switching frequencies, which are resolved by late-stage maturation into specific cell-types (*43, 44*). Our single-cell *in silico* lineage inference coupled with *in situ* HCR validation and single-cell multiplexed RNA imaging captured the timing, speed and maturation of secretory cell-types and their transcriptional programme across early epithelial progenitor lineages, while multiciliated progenitors originated from pluripotent cells and underwent maturation in a separate branch. Notably, we find early epithelial progenitors also present across *Xenopus tropicalis* non-neural ectoderm cells, expressing both goblet cell and ionocyte signatures. The *Xenopus tropicalis* developmental atlas spans all germ layers sampling many more cells, while this study is focused on the sampling of developing mucociliary epithelium and its cell-types. Within the double positive secretory cell-types at neurula stages, the goblet markers were expressed at higher mean levels (relative to ionocyte markers) and likely to have confounded the annotation and absence of early epithelial progenitors. The presented lineage inference uses expression data alone, without additional priors and assumes unidirectional progression without trans/de-differentiation. Future versions could incorporate spatial positioning, cell-shape dynamics, and cell cycle, including growth and division rates. Our data and lineage inference provide a quantitative and generalizable approach to characterising a continuous model of cell fate choice suited for temporal data, uncovering general features of embryonic MCE development.

We also find the type I and type II ionocytes that are functional equivalents of human alpha and beta submucosal epithelial serous acinar cells controlling ion flow, regulating fluid, bicarbonate secretion to hydrate mucins, balancing acid-base homeostasis and thereby the volume and composition of the air-surface liquid. Although human primary models (Air-liquid interface, human airway epithelial culture) are valuable, they are technically demanding and do not fully recapitulate relevant cell-type interactions occurring *in vivo*. Several MCE cell-types and hallmarks are also found in other tissues, including the pronephric kidney, gut, epididymis, and nasal cavity across species, with conserved TF families and signalling pathways. It is tempting to speculate that similar paradigms and gene regulatory networks exist across other MCE tissues to drive cell fate decisions. Through comparative analysis across 9 single-cell datasets, we identified conserved transcriptional programmes driving ciliogenesis in cell-types across vertebrates, while specialised cell-types, their cellular functions drive specific transcriptional programs across secretory and basal cell-types. We infer the regulatory relationships during MCE development through scRNA-seq, however future studies that apply single-cell multi-omic approaches (i.e., chromatin accessibility, transcriptomics, spatial positioning, etc.) will help integrate signalling regulation, transcription factor-enhancer crosstalk, further to push an improved understanding of cell fate decisions.

The current study provides novel insights into MCE development, however, there are some important considerations of note. Firstly, we leverage *Xenopus* animal cap organoids that mimic embryonic MCE development but have distinct commonalities and differences with the embryo. Secondly, while the highly abundant transcripts are robustly captured in the scRNA-seq data, many of the lowly expressed transcription factors, signalling components are not missed in the current analysis, and further lost during ortholog mapping to human ids. Thirdly, we observe expression of *foxa1*, a canonical marker of serotonin secretory cells, within single-cells at stage 22. These cells lack expression of other serotonin secretory markers, which is likely due to limited sequencing depth, but require further characterization to appreciate contribution to later developmental stages. Lastly, the insights from scRNA-seq analysis in this study are supported by *in situ* HCR and multiplex spatial RNA imaging. We hope that further functional studies will help understand mechanistic insights into the development and species-specific differences across mucociliary epithelia.

Our work provides a resource of cell state changes and developmental transitions accompanying MCE development and contributes to dissecting developmental mechanisms involved in the formation of mucociliary tissues. The identified and conserved molecular targets, mechanisms and cell-fate paradigms controlling MCE differentiation would be useful in the development of regenerative therapies, including ciliopathies, and chronic lung diseases, aimed at restoring mucociliary clearance.

## Acknowledgments

The authors thank Can Aztekin for in-situ HCR guidance; Helen Neil, Magali Michaut (reNEW Genomic Platform), Mie Mechta and the Single-Cell Omics Platform at the Novo Nordisk Foundation Center for Basic Metabolic Research (CBMR), and Jutta Bulkescher (reNEW Imaging Platform) for technical expertise, support and help. We also thank members of the Sedzinski, Kwon and Natarajan groups. We thank DeIC National HPC Centre (ABACUS 2.0 and eScience Center) and for computational resources. We also thank John B. Wallingford for discussions and comments on the manuscript.

## Author contributions

J.S., J.L., A.B. collected tissue samples and performed scRNA-seq; T.P. contributed to sequencing libraries; J.S., A.B., performed in-situ HCR; J.S., A.B., T.K., Y.K., H.S.L., T.J.P. contributed towards experimental validation; A.F.M., K.N.N. processed and analysed all data; S.C. and T.K. also contributed to data analysis; K.N.N wrote the manuscript with input from co-corresponding authors; J.S., T.K, and K.N.N. supervised the project.

## Data and materials availability

Sequencing data and processed gene counts are available on GEO (accession number GSE158088). The analysis scripts are available at https://github.com/Natarajanlab/XenopusMCEAtlas. The single-cell transcriptomics data and metadata can be downloaded for offline visualization or interactively visualized at CellxGene or UCSC cell browser.

## MATERIALS AND METHODS

### *Xenopus laevis* husbandry, embryos isolation and manipulation

Wild-type *Xenopus laevis* were obtained from Nasco-Wisconsin, Fort Atkinson, WI, USA. Frogs were maintained by centralised facilities according to common procedures provided by the international *Xenopus* community including National Xenopus Resource (NXR) at the Marine Biological Laboratory, Woods Hole, USA, the European Xenopus Resource Center (EXRC) at the University of Portsmouth, School of Biological Sciences, UK, and also Xenbase (http://xenbase.org). *Xenopus laevis* embryos were isolated and manipulated according to standard procedures (*68*): *Xenopus laevis* females were ovulated by injection of 500 U of human chorionic gonadotropin (HCG). The following day, eggs were collected, *in vitro* fertilised and dejellied in 2% cysteine (pH 7.9). Embryos were washed and reared in 1/3X Marc’s Modified Ringer’s (MMR) solution. Embryos were incubated until stage 8 according to Nieuwkoop and Faber (NF) developmental tables (*29*).

### *In vitro* animal cap culture, single-cell dissociation, library preparation and sequencing

A total of 100 *Xenopus laevis* animal caps from at least 3 independent embryo clutches were manually dissected from embryos at NF stage 8 as described in (*69*) and subsequently cultured in Danilchik’s for Amy (DFA) medium supplemented with 50 µg/ml gentamycin until desired developmental stage assessed by comparing to sibling embryos. From each of the 10 sampled embryonic stages (NF Stage 8, 10.5, 12, 13, 16, 18, 20, 22, 24 and 27), animal cap organoids were dissociated into a single-cell suspension by first washing with 0.01% BSA-PBS, then incubating with Newport buffer 2.0 described in (*24*) in a BSA (1 mg/ml) pretreated 1.5 ml Eppendorf at room temperature while agitating on thermomixer; typically about 5 min at 500 RPM for animal caps NF stages 8 to 22 and additional 10 min at 1200 RPM for NF stages 24 and 27. The single-cell suspension was spun down and washed with 0.01

% BSA-PBS twice at 300 rcf for 3 min, then spun down at 300 rcf for 5 min and resuspended in 0.01% BSA-PBS. Cells were passed through a 50 µm diameter cell strainer, and the concentration was adjusted to 500 cells/µl. We evaluated the quality and purity of single-cell in suspension by microscopy by staining with 2 µg/ml propidium iodide (PI) and 20 µm Hoechst 33342, and found negligible cell death (<1 %) up to one-hour post animal caps organoids dissociation. The single-cell suspension was left on ice for 15 min, before processing and loading using a wide-bore tip on the 10x chromium chip for droplet-based scRNA-seq v2 (stages 8, 13, 16, 18, 20, 24, 27) and v3 (stages 10.5, 12, 22), following manufacturers recommended protocol (*70*), and sequenced using paired-end reads on an Illumina NextSeq 500 (High output reagent kit v2.5 75 cycles) for 90 cycles with read 1 (26 bp, 10x bead barcode and UMIs), read 2 (56 bp, transcript) and index read (8 bp, 10x sample index).

### Whole-mount *in situ* hybridization and whole-mount *in situ* chain reaction (*in situ* HCR) and imaging

The *Xenopus laevis* embryos were *in situ* hybridised with *Cftr.L probes* according to (*71*). For *in situ* HCR on whole *Xenopus laevis* embryos were prepared as for a traditional *in situ* hybridization, up to the probe hybridization step, as described in (*71*)followed by the *in situ* HCR protocol described in (*60*). Hybridization probes were designed and generated by Molecular Instruments Inc. Embryos were fixed with 4% formaldehyde in 1X MEMFA salts in Wheaton vials for 60 minutes on a rotator at RT, washed 3X 15 minutes with PBS, permeabilized with 100% methanol overnight at -20°C. Following day, embryos were rehydrated with 5 minutes washed in: 1) 75% methanol / 25% H_2_0, 2) 50% methanol / 50% H_2_0, 3) 25% methanol / 75% H_2_0, then washed 2X 5 minutes in 100% PBST. Next, embryos were permeabilized with Proteinase K (final concentration 5 µg/ml in PBST) for 10 minutes at RT. Embryos were washed 3X 5 minutes in PBST to remove Proteinase K, washed 2X 5 minutes in 0.1 M triethanolamine. Following, 10 µl of acetic acid anhydride was added to the 0.1 M triethanolamine and embryos were incubated for 10 minutes at RT. Embryos were washed 2X 5 minutes with PBST and re-fixed in 4% formaldehyde in H_2_0 for 20 minutes at RT on a rotator. Next, embryos were washed 5X for 5 minutes with PBST. Embryos were washed 5 minutes in a preheated (37°C) 500 µl wash buffer (Molecular Instruments Inc.) and transferred to 1.5 ml Eppendorf tubes (10 embryos per tube). Washed buffer was removed and replaced by a preheated 500 µl hybridization buffer (Molecular Instruments Inc.) for a 30 minutes incubation at 37°C. The hybridization buffer was removed and replaced by a 500 µl probe solution prepared by diluting probe stock in the hybridization buffer to the final concentration of 6 nM). Subsequently, embryos were incubated at 37°C overnight, washed 2X 30 minutes with a wash buffer (Molecular Instruments Inc.), 5 minutes 50% 5XSSCT / 50% wash buffer for 5 minutes and 2X 20 minutes with 5X SSCT at RT. Embryos were preamplified with 1 ml amplification buffer (Molecular Instruments Inc.) for 10 minutes at RT. To visualise the probes, the preamplification buffer was replaced by an amplification solution prepared by first heating to 95°C the fluorescently-tagged hairpins pairs (h1 and h2 hairpins) corresponding to the designed probes. Hairpins were incubated for 30 minutes at RT in dark and subsequently added to 500 µl of amplification buffer (Molecular Instruments Inc.) and incubated for 5 minutes in dark. The final concentration of hairpins was 48 nM. Preamplification buffer was removed from embryos and replaced by the hairpin solution. Embryos were incubated overnight at RT in the dark. The following day, embryos were transferred to Wheaton vials and washed 2X with 5X SSCT for 30 minutes and stored in 1X PBS at 4°C. For immunostaining after *in situ*, embryos were blocked in 1ml 50% Cas-Block / 50% PBST for 1 hour in Wheaton vials at RT in dark followed by incubation with the primary antibody (anti-Acetylated-tubulin, Sigma, T7451, 1:500) in Cas-Block at 4°C overnight in dark. The next day, embryos were washed 3X 10 minutes with PBST and incubated with a secondary antibody (anti-mouse-647, Thermo Fisher, A21235, 1:250) diluted in 1 ml 50% Cas-Block / 50% PBST for 1 hour at RT. Embryos were washed 3X for 10 minutes with PBST in dark and mounted as described in (*72*) imaged using Zeiss 880 inverted confocal microscope using C-Apochromat 40x/1.2 W Corr M27 objective. For sectioning, embryos were embedded in 2% low-melting-point agarose and sectioned using a vibratome set up to a 50 µm section thickness, stained with 0.1 µg/ml DAPI (Thermo Fisher, D1306) in 1X PBS at RT for 10 minutes, mounted in Moviol (10% Mowiol 4-88, 25% glycerol, 0.1 M Tris/pH 8.5, 2% N-propyl gallate), with coverslip and imaged using Zeiss 880 inverted confocal microscope using C-Apochromat 40x/1.2 W Corr M27 objective. Fiji (*73*) was used for image processing including maximum intensity projection of acquired z-stacks and contrast adjustment.

### Molecular cartography

#### Sample fixation

*Xenopus laevis* embryos were fixed and stabilised according to Resolve guidelines. Briefly, embryos were incubated at room temperature in PAXgene^Ⓡ^ fixation solution for 4 hours followed by 48 hours of incubation at room temperature in PAXgene^Ⓡ^stabiliser. Next, embryos were incubated for 48 hours in 20% Sucrose (w/v in 1X PBS) at 4℃ until embryos sank to the bottom of the well.

#### Cryo-embedding and cryo-sectioning

Embryos were washed of 20% Sucrose (w/v in 1X PBS) in 1X PBS/OCT (optimal cutting temperature medium, Neg-50 Frozen Section Medium, VWR) (2:1, 1:1, 1:2, 100% OCT), 30 minutes each wash at room temperature. Next, embryos were placed in the middle of the cryo-embedding medium. The cryomold was placed on top of the liquid nitrogen in the gas phase until the medium was completely white. The cryo-embedded block was kept in dry ice until sectioning. Embryos were sections at 12 µm and sections were collected onto a precooled coverslip. The samples were sent to Resolve GmbH, Germany for their processing.

#### Mol. Cartography

PAXgene fixed samples were used for Molecular Cartography™ according to the manufacturer’s instructions (protocol 3.0; available for download from Resolve’s website to registered users), starting with the aspiration of ethanol and the addition of buffer BST1 (step 6 and 7 of the tissue priming protocol). Briefly, tissues were primed followed by overnight hybridization of all probes specific for the target genes (see below for probe design details and target list). Samples were washed the next day to remove excess probes and fluorescently tagged in a two-step colour development process. Regions of interest were imaged as described below and fluorescent signals were removed during decolourization. Colour development, imaging and decolourization were repeated for multiple cycles to build a unique combinatorial code for every target gene that was derived from raw images as described below.

#### Probe design

The probes for 98 genes were designed using Resolve’s proprietary design algorithm. Briefly, the probe design was performed at the gene level. For every targeted gene all full-length protein-coding transcript sequences from the ENSEMBL database were used as design targets if the isoform had the GENCODE annotation tag ‘basic’ (*74, 75*). To speed up the process, the calculation of computationally expensive parts, especially the off-target searches, the selection of probe sequences was not performed randomly, but limited to sequences with high success rates. To filter highly repetitive regions, the abundance of *k*-mers was obtained from the background transcriptome using *Jellyfish* (*76*). Every target sequence was scanned once for all *k*-mers, and those regions with rare *k*- mers were preferred as seeds for full probe design. A probe candidate was generated by extending a seed sequence until a certain target stability was reached. A set of simple rules was applied to discard sequences that were found experimentally to cause problems. After these fast screens, every kept probe candidate was mapped to the background transcriptome using *ThermonucleotideBLAST* (*76*) and probes with stable off-target hits were discarded. Specific probes were then scored based on the number of on-target matches (isoforms), which were weighted by their associated APPRIS level (*77*), favouring principal isoforms over others. A bonus was added if the binding-site was inside the protein-coding region. From the pool of accepted probes, the final set was composed by greedily picking the highest-scoring probes.

#### Imaging

Samples were imaged on a Zeiss Celldiscoverer 7, using the 50x Plan Apochromat water immersion objective with an NA of 1.2 and the 0.5x magnification changer, resulting in a 25x final magnification. Standard CD7 LED excitation light source, filters, and dichroic mirrors were used together with customised emission filters optimised for detecting specific signals. Excitation time per image was 1000 ms for each channel (DAPI was 20 ms). A z-stack was taken at each region with a distance per z-slice according to the Nyquist-Shannon sampling theorem. The custom CD7 CMOS camera (Zeiss Axiocam Mono 712, 3.45 µm pixel size) was used. For each region, a z-stack per fluorescent colour (two colours) was imaged per imaging round. A total of eight imaging rounds were done for each position, resulting in 16 z-stacks per region. The completely automated imaging process per round (including water immersion generation and precise relocation of regions to image in all three dimensions) was realised by a custom python script using the scripting API of the Zeiss ZEN software (Open application development).

#### Spot segmentation

The algorithms for spot segmentation were written in Java and are based on the ImageJ library functionalities. Only the iterative closest point algorithm is written in C++ based on the libpointmatcher library (https://github.com/ethz-asl/libpointmatcher).

#### Preprocessing

As a first step all images were corrected for background fluorescence. A target value for the allowed number of maxima was determined based upon the area of the slice in µm² multiplied by the factor 0.5. This factor was empirically optimised. The brightest maxima per plane were determined, based upon an empirically optimised threshold. The number and location of the respective maxima was stored. This procedure was done for every image slice independently. Maxima that did not have a neighbouring maximum in an adjacent slice (called z-group) were excluded. The resulting maxima list was further filtered in an iterative loop by adjusting the allowed thresholds for (Babs-Bback) and (Bperi-Bback) to reach a feature target value (Babs: absolute brightness, Bback: local background, Bperi: background of periphery within 1 pixel). This feature target values were based upon the volume of the 3D-image. Only maxima still in a z-group of at least 2 after filtering were passing the filter step. Each z-group was counted as one hit. The members of the z-groups with the highest absolute brightness were used as features and written to a file. They resemble a 3D-point cloud. *Final segmentation and decoding:* To align the raw data images from different imaging rounds, images had to be corrected. To do so the extracted feature point clouds were used to find the transformation matrices. For this purpose, an iterative closest point cloud algorithm was used to minimise the error between two point-clouds. The point clouds of each round were aligned to the point cloud of round one (reference point cloud). The corresponding point clouds were stored for downstream processes. Based upon the transformation matrices the corresponding images were processed by a rigid transformation using trilinear interpolation.The aligned images were used to create a profile for each pixel consisting of 16 values (16 images from two colour channels in 8 imaging rounds). The pixel profiles were filtered for a variance from zero normalised by the total brightness of all pixels in the profile. Matched pixel profiles with the highest score were assigned as an ID to the pixel. Pixels with neighbours having the same ID were grouped. The pixel groups were filtered by group size, number of direct adjacent pixels in the group, and number of dimensions with a size of two pixels. The local 3D-maxima of the groups were determined as potential final transcript locations. Maxima were filtered by the number of maxima in the raw data images where a maximum was expected. The remaining maxima were further evaluated by the fit to the corresponding code. The remaining maxima were written to the results file and considered to resemble transcripts of the corresponding gene. The ratio of signals matching to codes used in the experiment and signals matching to codes not used in the experiment were used as estimation for specificity (false positives).

### scRNA-seq: Data processing (alignment and quality control)

The raw 10x genomics sequencing data were processed using the CellRanger (version 2.1.1) with the default option for 3’ Gene Expression v2 Libraries (‘SC3Pv2’). The sequences were mapped to *Xenopus laevis* genome (XenBase RRID# SCR_003280; NCBI GCA_001663975.1) using annotation from the GitLab repository (https://gitlab.com/Xenbase/genomics/XENLA_9.2; 2018-May version) (*49, 78*). The 10 individual scRNA-seq datasets were aligned to (XENLA_GCA001663975v1_XBv9p2) reference genome using zUMI’s alignment and feature counting (version 2.4.5b), which utilises STAR (version 2.5.4b) (*79, 80*). The XENLA 9.2 assembly included two annotations for gene otogelin (*mucXS*) referred as *Otog* and *otogl2.* To filter out background barcodes we applied EmptyDrops (from package DropletUtils version 1.2.2) using default parameters and selected only cells passing a < 0.01 FDR threshold, for every individual dataset (*81*). To identify doublets we applied scrublet (version 0.2.1) using parameters expected_doublet_rate = 0.06, min counts = 2, min_cells = 3, min_gene_variability_pctl = 85 and n_prin_comps = 30 to all datasets individually (*81, 82*). We normalized the datasets for cell library size, log-transformed and scaled genes to mean zero and unit variance, and identified highly variable genes using scanpy (version 1.4) (*83*).

### scRNA-seq: Data visualisation and marker gene detection

For each MCE developmental stage, we performed dimensionality reduction using UMAP (*84*) with 10 neighbours on 40 principal components and identified clusters using Louvain community detection (*85*). To generate an integrated force-directed knn-graph layout, we applied SPRING to the integrated single-cell expression data, using default parameters (*86*). We performed a Wilcoxon rank-sum test between each identified cluster and the remaining cells in the integrated dataset using scanpy function scanpy.tl.rank_genes_groups with use_raw = True.

#### Imputation

We imputed gene counts using MAGIC (*86*) (using palantir.utils.run_magic_imputation using parameters n_steps = 3. *Note: Imputed values were only used for visualisation of marker gene expression of embeddings and to infer gene trends*.

#### Developmental dynamics

To infer pseudotime and developmental potential we applied Palantir (version 0.2.1) using default parameters (*87*). To set the start cell we selected one arbitrary cell from the earliest sample timepoint. To run Palantir on discrete time-series data we “stitched” it together with harmony (*88*) (version 0.1.1) using harmony.core.augmented_affinity_matrix with parameters n_neighbors = 20 and pc_components = 50. Diffusion maps (also from the Palantir package) were computed with parameters n_components = 20, knn = 20. To validate the inferred developmental dynamics, we applied CytoTRACE (version 0.1.0) using the function iCytoTRACE for dataset integration with default parameters (*44*).

#### Phenograph

To cluster the integrated dataset we used Phenograph (version 1.5.2) (*86*), through Palantir function Palantir.utils.determine_cell_clusters with parameters k = 1000.

#### Louvain

Individual stages were clustered using louvain, through scanpy.tl.louvain, using 10 neighbours and 40 principal components. Louvain resolution parameters were set for each stage individually (in range 0.01 to 0.3).

#### Cluster stability

Robustness of clustering was assessed by repeating 20 independent iterations of Phenograph and louvain clustering for each developmental stage 20 times, using different random seeds and calculating element-wise clustering consistency (ECS; element_consistency) using ClustAssess package (version 0.3.1) with default parameters (*89*).

#### Gene trends

Gene trends over branches were computed using Palantir function palantir.presult.compute_gene_trends on the MAGIC imputed gene counts (*86*).

#### Gene Ontology

We use ClusterProfiler (version 3.10.1) for GO analysis (*90*). Xenopus.laevis gene names were mapped to human orthologs. We used enrichGO per stage to cluster gene sets, to assert significantly enriched biological processes. Comparison between individual stages or clusters were achieved using the compareCluster function. Significant terms were selected with a p < 0.05 threshold, after Benjamini-Hochberg correction for multiple testing.

#### Lineage inference

To infer the developmental lineages we applied a neighbour mapping and voting strategy. Here each cell votes for the cluster of its nearest neighbour at the preceding timepoint. Votes are aggregated within clusters and a link is drawn to the most likely developmental “ancestor”. Neighbour search is performed with Scipy (version 1.1.0) function scipy.spatial.cKDTree using default parameters. Upon assessment of voting confidence and marker gene expression, low confidence links were refined using the expression of key branch marker genes.

#### Gene set quantification

To quantify the utilisation of different functional gene groups, we plotted the normalised and z-scored gene expression of the cell cycle, transcription factor and signalling gene sets, retrieved from public databases.

#### Cell-cycle

Cell Cycle genes were retrieved from cyclebase (*91*).

#### Transcription factors

Human TF’s were retrieved from AnimalTFDB 3.0 (*41*).

#### Signalling

Signalling gene sets were obtained from msigdb (gsea-msigdb.org/) and converted to *X. laevis* orthologs. A handful of *X. laevis* gene sets were removed (*hvl, hdac* from Notch pathway, *smad2/3/4* from Wnt pathway, *rho, rock1/2, e2f4/5* from TGF-beta pathway) and manually added (*wnt, fzd, lrp5* to Wnt pathway, *tgfbr1, gdf11.1, nodal inhbc2* to TGF-beta pathway).

#### Cell cycle classification

The cell cycle stage prediction was performed using the scanpy function scanpy.tl.score_genes_cell_cycle) to score S and G2M phase genes. The reference cell cycle stage marker genes from (*92*) can be found here (https://github.com/theislab/scanpy_usage/blob/master/180209_cell_cycle/data/regev_lab_cell_cycle_g <u>enes.txt</u>). Each single cell is assigned an S and G2M score, and classified based on the highest-scoring gene set. If neither gene set scores above 0.5, the cell is classified as G1.

#### RNA velocity

On the zUMIs generated intron-exon classified output, we applied scVelo (version 0.2.2) (*39*) per stage and branch. scVelo were run on 10 principal components and 50 neighbours, in “stochastic” mode.

#### Mapping to bulk Xenopus datasets

Expression data were retrieved through GEO (GSE76342) (*91*). Gene names were mapped to *Xenopus laevis* gene names. If several genes map to a *Xenopus laevis* gene name, only the first is retained. To assess similarity we performed spearman correlation between the two datasets, on all HVG’s also detected in (*42*), totalling 1376 genes).

#### Mapping cell-types from Xenopus tropicalis single-cell atlas

The non-neural ectoderm lineage cells were re-analysed from Xenopus tropicalis single-cell developmental atlas (*24*) and compared to our cell-types. A gene set consisting of 33 goblet markers and 20 ionocyte markers was selected, which robustly marked the respective cell-types across both studies (based on author annotation). The mean expression of the respective goblet cell and ionocyte gene sets was used to re-classify single-cells as single positive (Gc^+^ or Ic^+^) or double-positive cells (Gc^+^, Ic^+^), from non-neural ectoderm lineage from all stages.

#### Comparative evolutionary development comparison

We downloaded eight single-cell datasets spanning *Xenopus*, mouse and human nasal/airway epithelium atlases from respective repositories, spanning 144 cell-types across 120,842 single-cells (Supplementary note 4). We converted all gene symbols to human orthologs and used them as a common reference to project datasets onto each other. For each dataset and respective cell-types, marker genes were identified first using scanpy (p < 0.05). All marker genes from all cell-types within each data were combined to form dataset-specific gene sets, and enrichment analysis (per cell-type) was calculated, *i.e.,* the mean expression in a given cell-type relative to the mean expression within the dataset. The individual cell-type gene enrichments were pairwise-correlated (Rank-based Spearman correlation) within and across datasets. To define the basal, secretory and multiciliated gene sets, we used differential expressed genes identified in at least 20% of cell-types within each cluster. Each cell-type was subsequently scored (*enrichment score*) on the basis of gene expression similarity to the conserved gene sets within the clusters of cell-types. The conserved marker genes (Fig. S15E) were identified by counting the number of times a given gene was classified as a marker, and ranked based on frequency, i.e., how often genes were classified as markers in either multiciliated, ionocyte or goblet cell types and across all datasets.

#### Multiplexed single-cell spatial analysis (Spot analysis, quantification and correlation)

The nuclear segmentation of cells was performed using manufacturer recommendations, *i.e.,* the maximum intensity projection using watershed cell detection in QuPath (version 0.2.3) with parameters (background radius = 25 px, median filter radius = 6 px, sigma = 8 px, minimum area = 25 px^2, maximum area = 6000 px^2, threshold = 30, cell expansion = 50 px and smooth boundaries enabled). Spots were quantified in each region of interest using Polylux version 1.6.1 (Resolve Biosciences) within Fiji/ImageJ2 version 2.1.0/1.53c. Individual segmented cells with less than 8 spot counts were disregarded, and spot counts were normalized to the same number of spots per cell, to facilitate comparison between cells.

Normalized spot counts (>8 spots) for marker genes probes were superimposed over DAPI segmented nuclei and sectioned images. The relative marker gene expression across stages was calculated for all cells passing the quality control, followed by single-cell classification into either multiciliated, goblet, basal cells or ionocytes. The heatmap was plotted using scaled mean expression across all stages.

