## Supplementary Figures for "A single-cell, time-resolved profiling of Xenopus mucociliary epithelium reveals non-hierarchical model of development"

### Supplementary figure 1

A

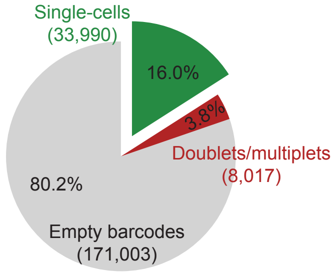


| Stages | #cells |
|---|---|
| 8 | 3408 |
| 10.5 | 1261 |
| 12 | 868 |
| 13 | 5823 |
| 16 | 3252 |
| 18 | 4228 |
| 20 | 3450 |
| 22 | 5222 |
| 24 | 3107 |
| 27 | 3371 |

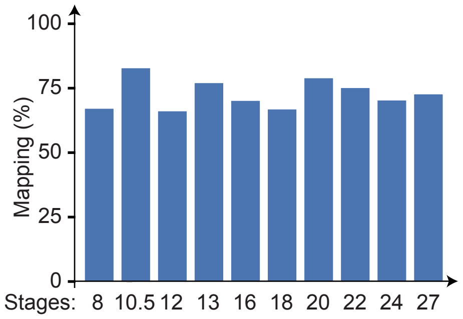


B

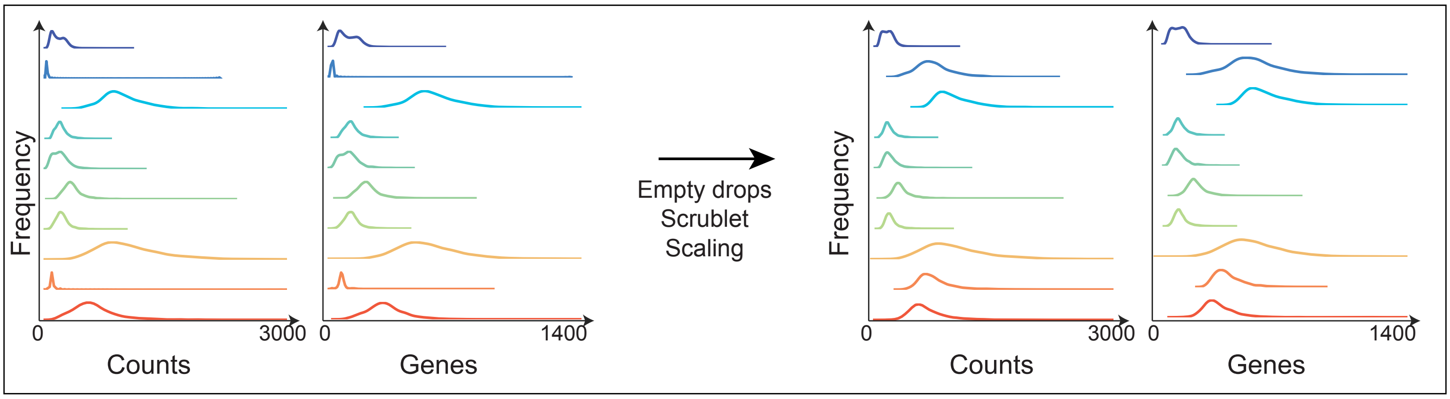


C

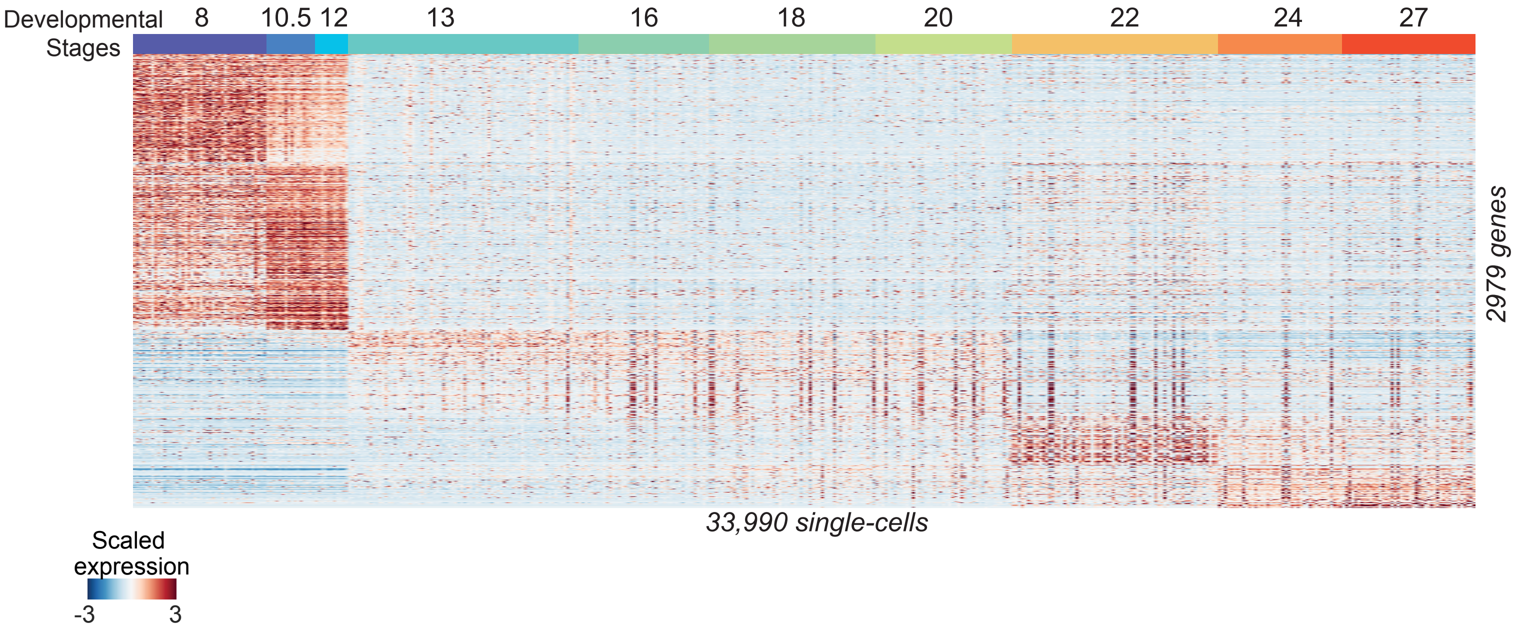

**Fig. S1**(**A**) Relative proportion of cells passing quality control after droplet scRNA-seq and mapping statistics across 10 stages of developing MCE.

(**B**) The expression distribution (counts and genes) before and after quality control for each developmental stage. The quality control analysis is performed to remove empty (residual/floating RNA), doublet/multiples and scale across different MCE stages.

(**C**) Expression pattern of 2979 highly variable genes (HVGs) over 33,990 single-cells across 10 developmental stages.

### Supplementary figure 2

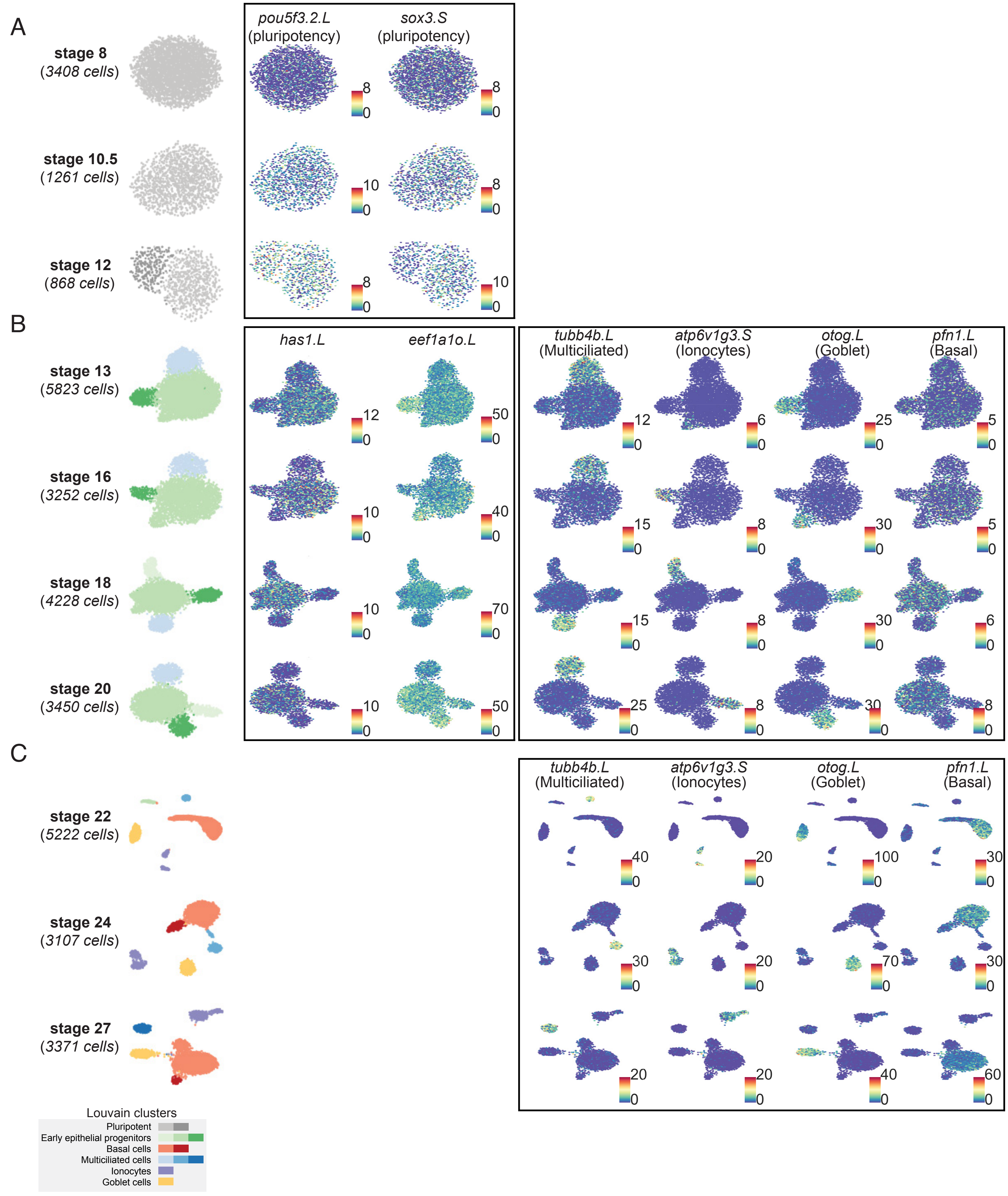

**Fig. S2**(**A**) UMAP plots showing blastula (stage 8) and gastrula stage (NF 10 and 12.5) cells marked by specific expression of pluripotency factors (*pou5f3.2.L* and *sox3.S*).

(**B**) Similar to **(A)**, but during neurula stages marking progenitor populations of multiciliated (*tubb4b.L*), ionocytes (*atp6v1g3.S*), goblet (*otog.L*) and basal cells (*pfn1.L*). The carbohydrate polymer hyaluronan synthase (*has1.L*) and elongation factor (*eef1a1o.L*) are expressed throughout neurula stages in all progenitors.

(**C**) Similar to **(A, B)**, but during early tailbud stages marking terminal cell types including multiciliated (*tubb4b.L*), ionocytes (*atp6v1g3.S*), goblet (*otog.L*) and basal (*pfn1.L*) cells.

The cells (left) are colored by cell types identified by Louvain clustering. The scale bars indicate relative z-scaled expression across cells.

### Supplementary figure 3

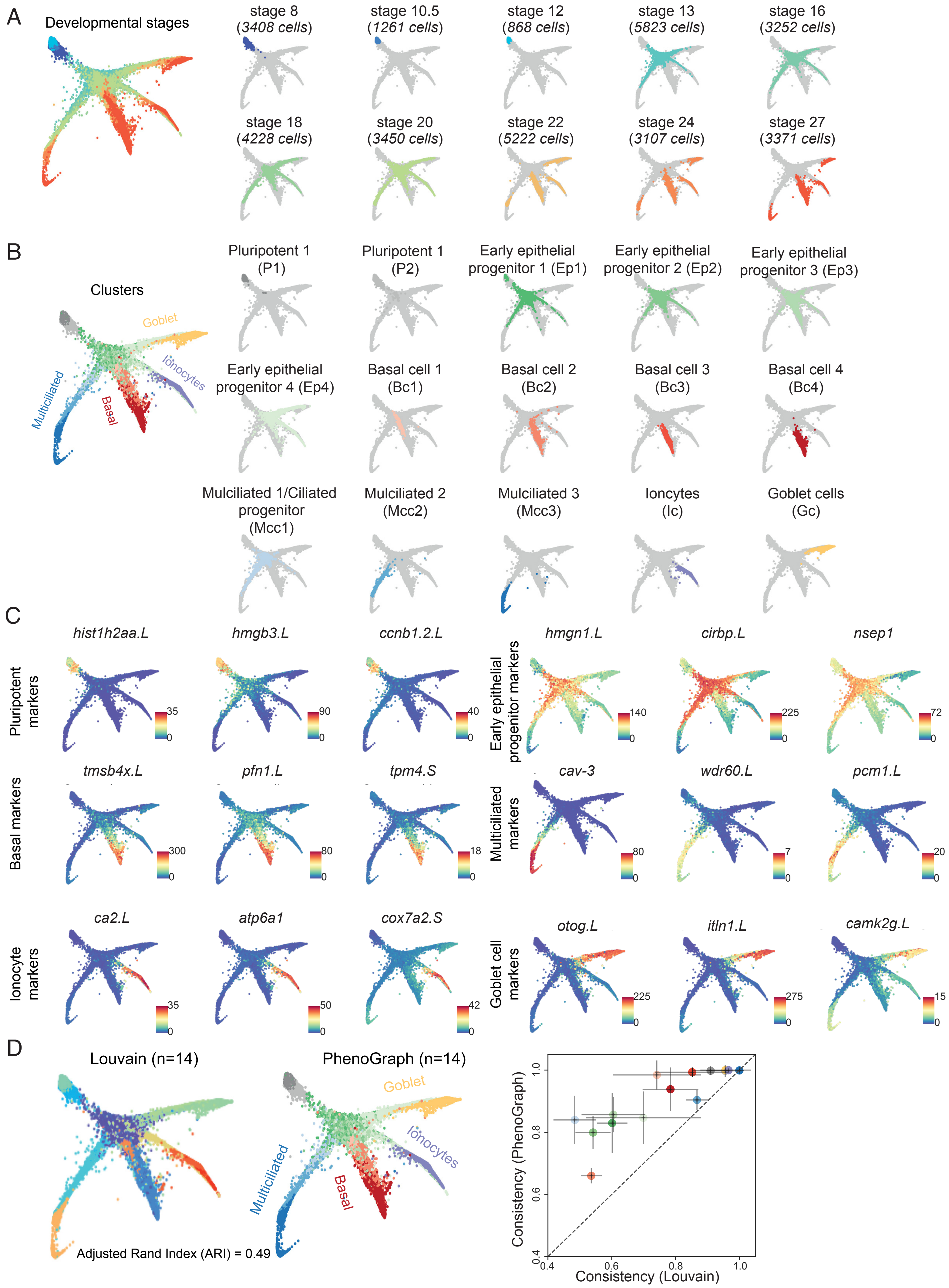

**Fig. S3**(**A**) Embedding of single-cells during MCE development (MCE manifold) over a knn graph colored by stages (left). The number of cells across individual stages and their positions are overlaid and colored on knn-graph (right).

(**B**) Phenograph clusters highlighted over the MCE developmental manifold (left). The PhenoGraph clusters and respective cells are overlaid and colored on knn-graph (right).

(**C**) Expression of marker genes over MCE developmental manifold. The scale bars indicated the scaled imputed expression of respective markers.

(**D**) Adjusted Rand Index (ARI) highlighting the dissimilarity of the identified clusters between PhenoGraph and community-based Louvain clustering. Louvain resolution was set to achieve the same number of clusters for comparison (n), for this comparison. ARI, a measure of cluster similarity, indicated only a 49% match between two clustering approaches. Consistency was computed using element-wise consistency (ECS), using random seeds and default resolution parameter.

Supplementary figure 4

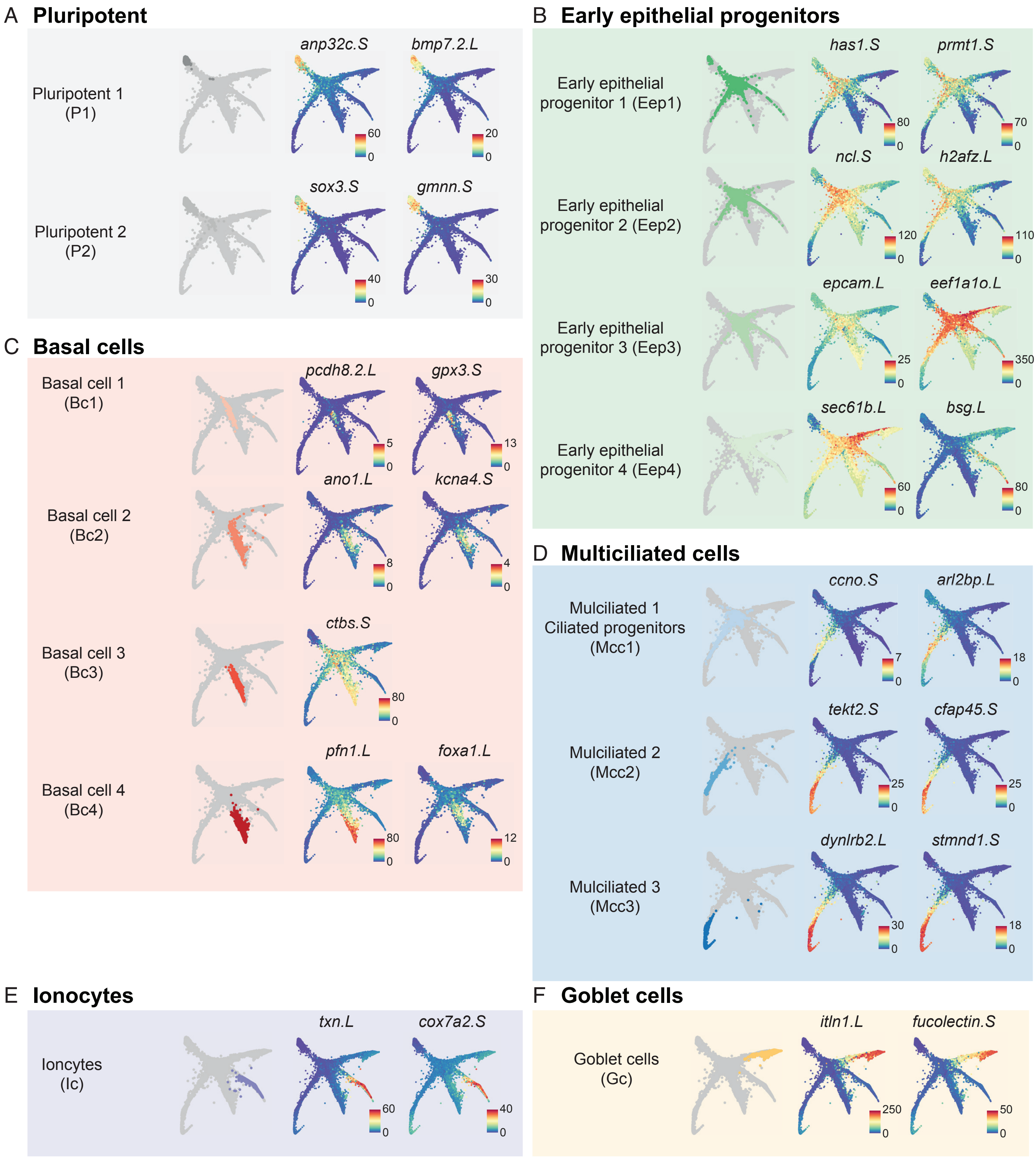

**Fig. S4** (**A**) MCE developmental knn graph showing expression of marker genes for each cell type and PhenoGraph cluster incl. pluripotent cells (A), early epithelial progenitors **(B)**, basal **(C)**, multiciliated **(D)**, ionocytes **(E)** and goblet cells **(F)**.

The scale bars indicated the scaled imputed expression of respective markers.

Supplementary figure 5

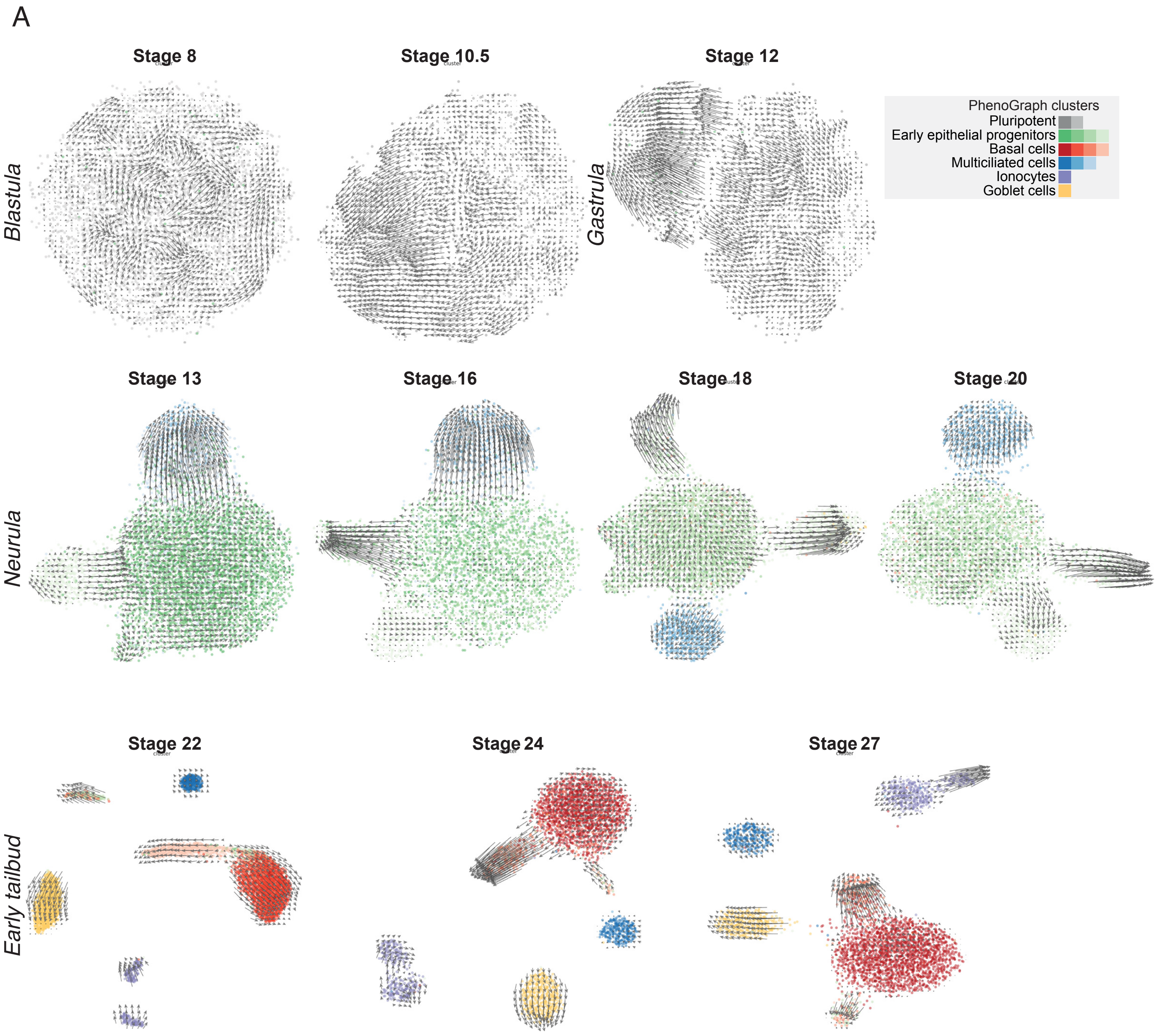

**Fig. S5**(**A**) RNA velocity inferred over each MCE developmental stage and visualised over UMAP to highlight developmental propensities.

Supplementary figure 6

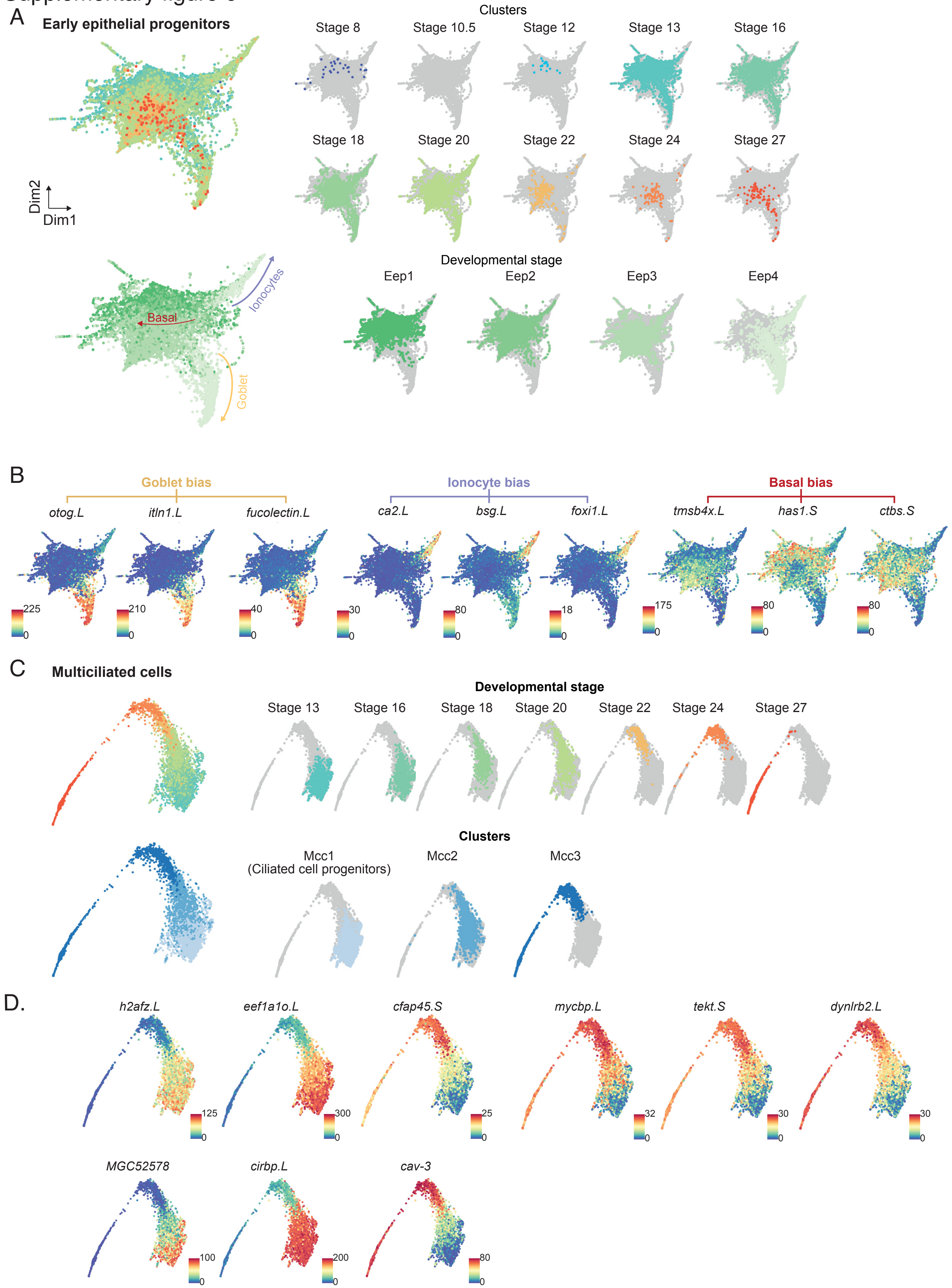

**Fig. S6**(**A**) Low dimensional graph embedding and visualisation of early epithelial progenitor cells (left), overlaid with cells from individual MCE developmental stages (top) and PhenoGraph clusters (bottom). The multi-lineage bias of early epithelial progenitors is indicated by arrows.

(**B**) Low dimensional visualisation of early epithelial progenitor cells indicating lineage bias, and overlaid with the expression of cell type markers including goblet (*otog.L*, *itln1.L*, *fucolectin.L*), ionocytes (*ca2.L*, *bsg.L*, *foxi1.L*) and basal cells (*tmsb4x.L*, *has1.S*, *ctbs.S*). The scale bars indicate the scaled imputed expression of respective markers.

(**C**) Low dimensional visualisation of multiciliated cells alone (left), overlaid with cells from individual MCE developmental stages (top) and PhenoGraph clusters (bottom).

(**D**) Low dimensional visualisation of multiciliated cells overlaid with specific markers (*h2afz.L*, *eef1a10.L*, *cfap45.S*, *mycbp.L*, *tekt.S*, *dynlrb2.L*, *MGC52578*, *cirbp.L*, *cav-3*). The scale bars indicate the scaled imputed expression of respective markers.

Supplementary figure 7

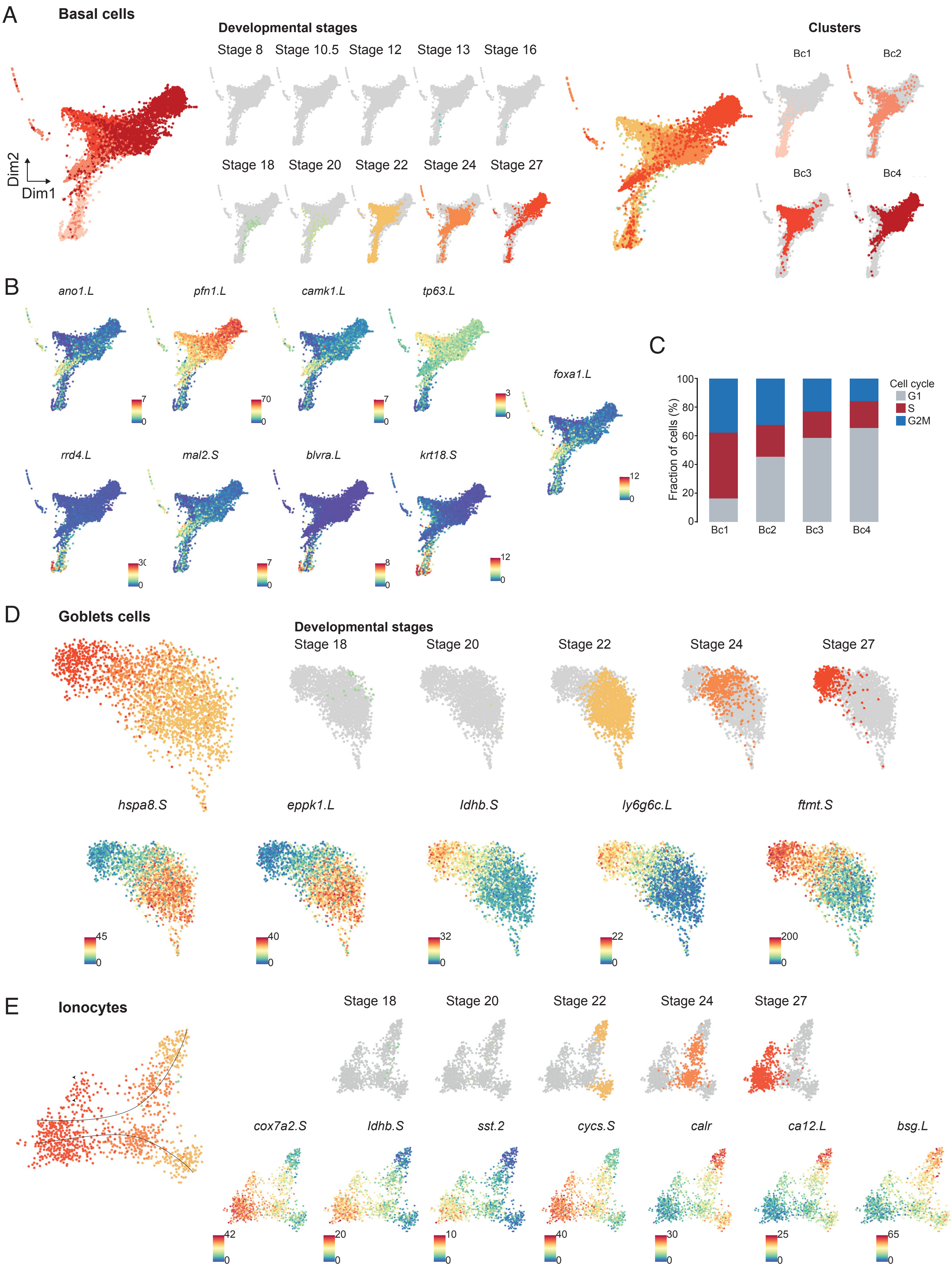

**Fig. S7**(**A**) Low dimensional visualisation of basal cells alone (left), overlaid with cells from individual MCE developmental stages (top) and PhenoGraph clusters (bottom).

(**B**) Low dimensional visualisation of basal cells overlaid with specific markers (*ano1.L*, *pfn1.L*, *camk1.L*, *rrd4.L*, *mal2.S*, *blvra.L*, *tp63.L*, *krt18.S*), including small secretory cell marker (SSC: *foxa1.L*).

The scale bars indicate the scaled imputed expression of respective markers.

(**C**) The basal cell subclusters can also be differentiated based on the proportion of cells in different cell cycle stages.

(**D**) Low dimensional visualisation of goblet cells alone (left), overlaid with cells from individual MCE developmental stages (top) and overlaid with specific markers (*hspa8.S*, *tmsb4x.L*, *ly6g6c.L*, *Idhb.S*, *eppk1.L*, *ftmt.S*).

(**E**) Low dimensional visualisation of ionocyte cell subpopulations alone (left), overlaid with cells from individual MCE developmental stages (top) and overlaid with specific markers (*tmsb4x.L*, *cox7a2.S*, *ca12.L*, *cycs.S*, *calr*, *Idhb.S*, *sst2*, *bsg.L*). Two ionocyte subtypes can be observed based on marker expression over developmental stages (black arrows)

Supplementary figure 8

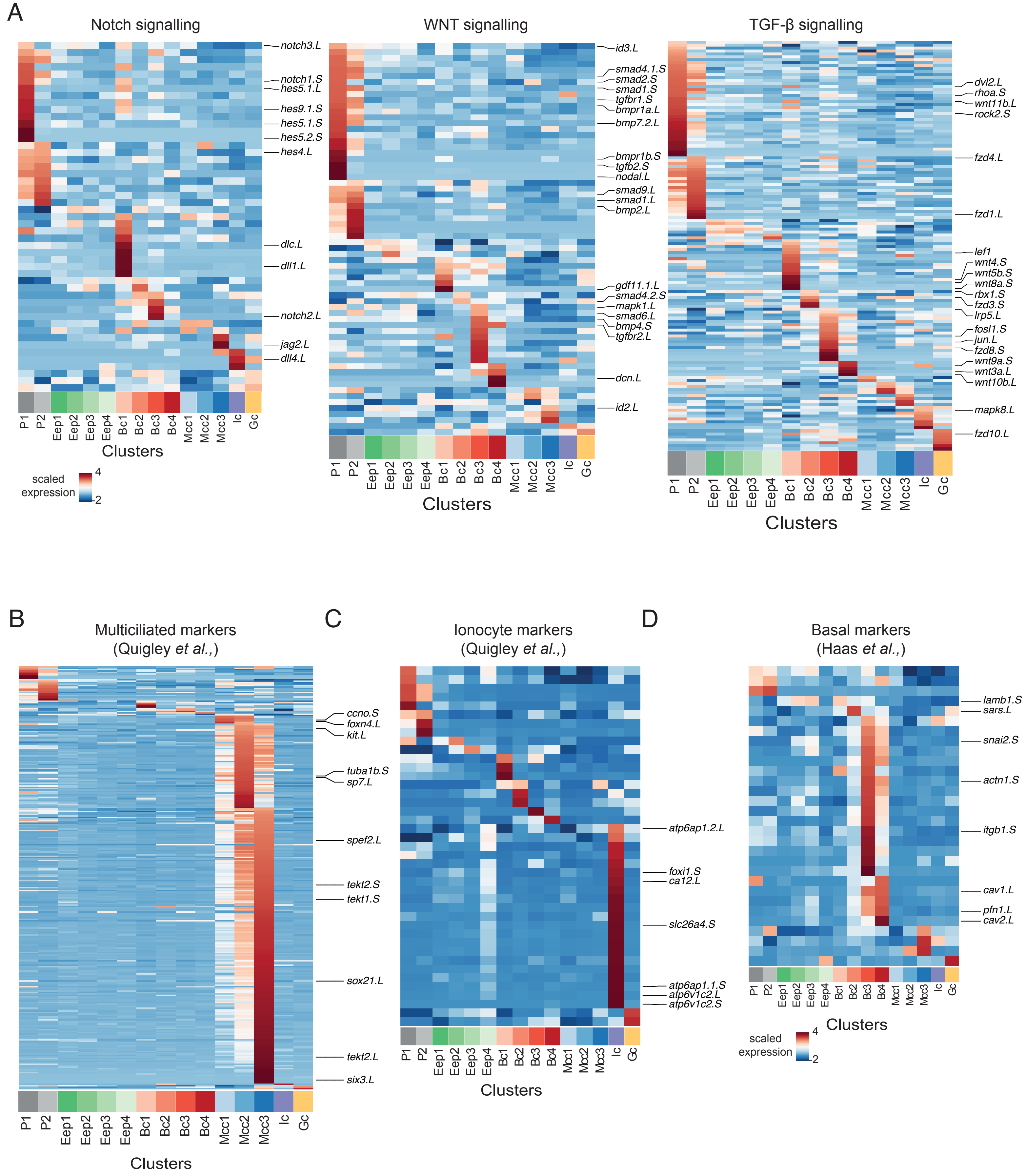

**Fig. S8**(**A**) Heatmap showing the expression pattern of Notch, Wnt and TGF-B signalling pathways over MCE developmental stages and PhenoGraph clusters respectively.

(**B**) Heatmap showing the expression pattern of multiciliated core genes (y-axis), identified across MCE PhenoGraph clusters (x-axis). Step-wise expression and maturation of MCC core genes can be observed from Mcc1 to Mcc2 to Mcc3. While most multiciliated markers are indeed expressed in MCC subclusters, many bulk genes are heterogeneously expressed in basal, goblet cells and ionocytes.

(**C**) Heatmap showing the expression pattern of ionocyte core genes (y-axis) over MCE PhenoGraph clusters (x-axis).

(**D**) Heatmap showing the expression pattern of basal cells core genes (y-axis) over MCE PhenoGraph clusters (x-axis).

The scale bars indicate the z-scaled mean expression of respective markers.

### Supplementary figure 9

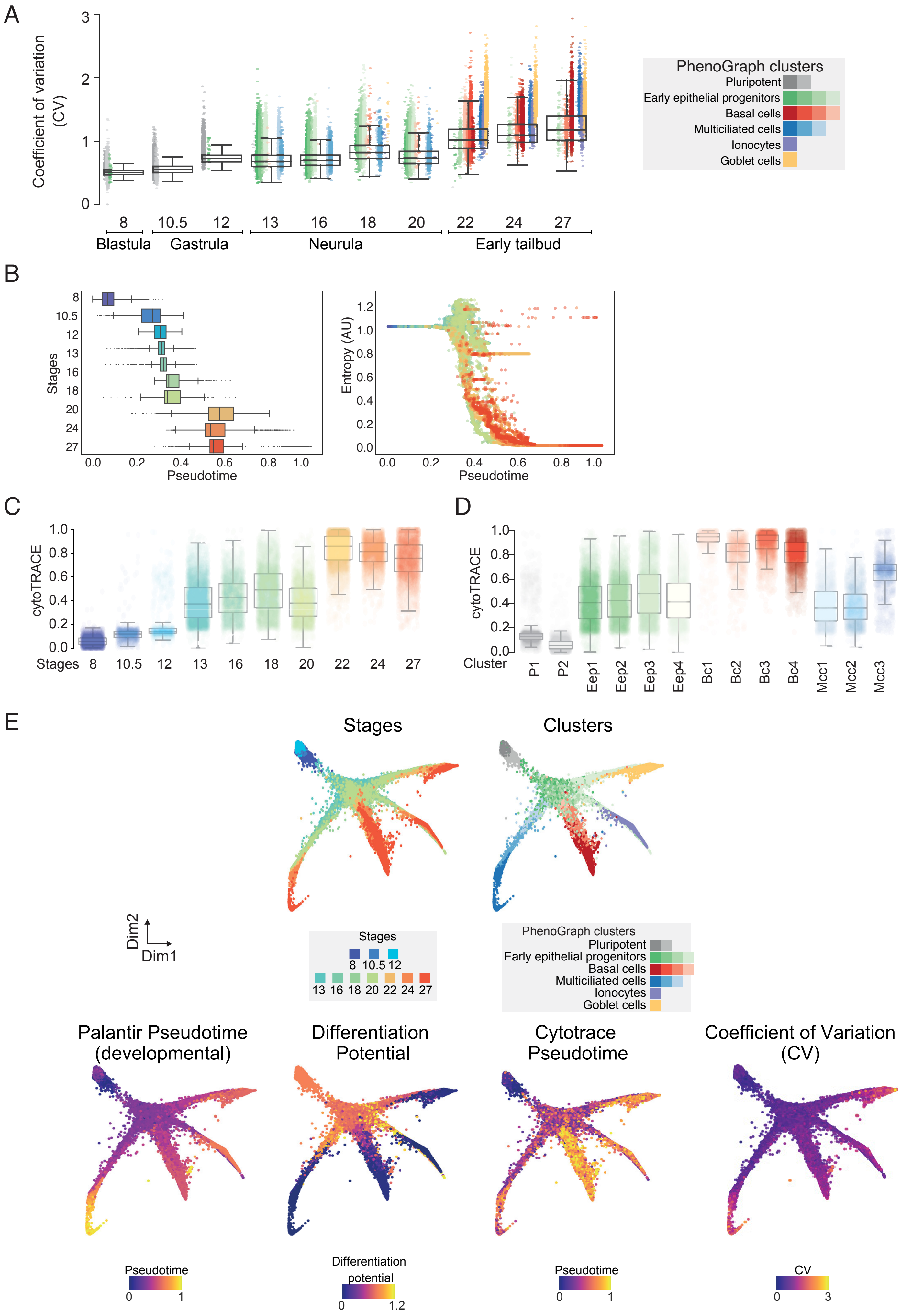

**Fig. S9**(**A**) Boxplots comparing MCE developmental stages (x-axis) and subclusters (colored points) with the coefficient of variation (CV) for MCE PhenoGraph clusters. The CV increases over development (highest across early progenitors and basal cells), indicating heterogeneity and stochasticity as a hallmark to drive noise-induced differentiation.

(**B**) Boxplots comparing MCE developmental stages (left) and single-cell entropy (right) with developmental pseudotime ordering (x-axis).

(**C**) Boxplots comparing MCE developmental stages (x-axis) with cytoTrace pseudotime scores.

(**D**) Boxplots comparing PhenoGraph clusters (x-axis) with cytoTrace pseudotime scores.

The low cytoTrace scores (pluripotent, early epithelial and ciliated progenitors) indicate high stemness scores, while late-stage cell types have high cytoTrace scores indicating a differentiated state.

(**E**) Overview of different molecular features contributing to MCE developmental and cell-fate specification, including MCE developmental knn graph overlaid with MCE developmental stages, PhenoGraph clusters, Palantir developmental Pseudotime, Differentiation potential, CytoTrace Pseudotime and Coefficient of variation (CV).

Supplementary figure 10

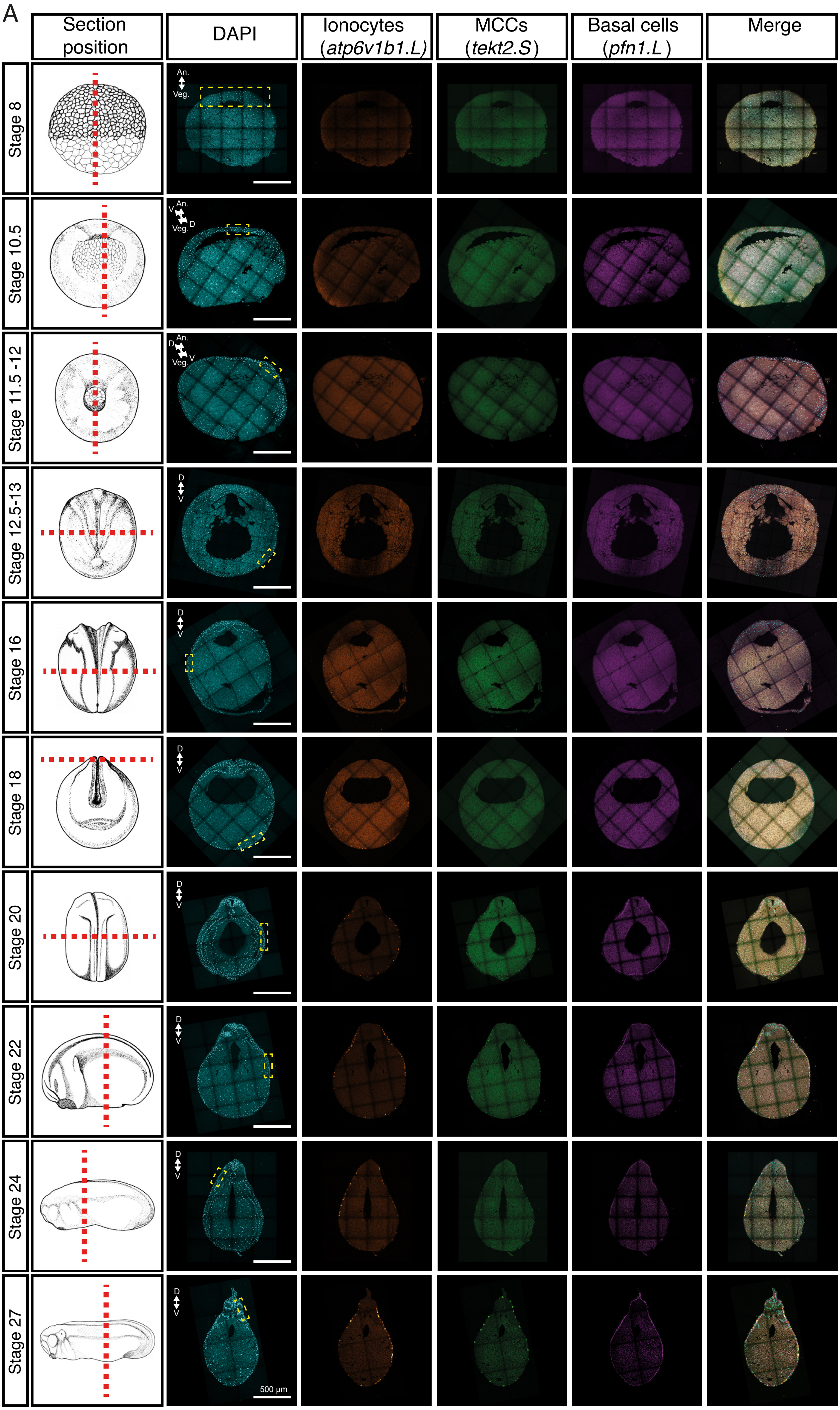

**Fig. S10**(**A**) *in-situ* hybridization chain reaction (HCR) and validation of lineage inference over 10 development stages of the embryonic epidermis in embryos, marking ionocytes (*atp6v1b1.L*, orange), multiciliated (*tekt2.S*, green) and basal cells (*pfn1.L*, magenta). The nuclei are marked by DAPI staining (cyan). The yellow rectangles indicate zoomed-in regions shown in Figure 6A. Images represent maximum intensity projections of transverse sections.

Supplementary figure 11

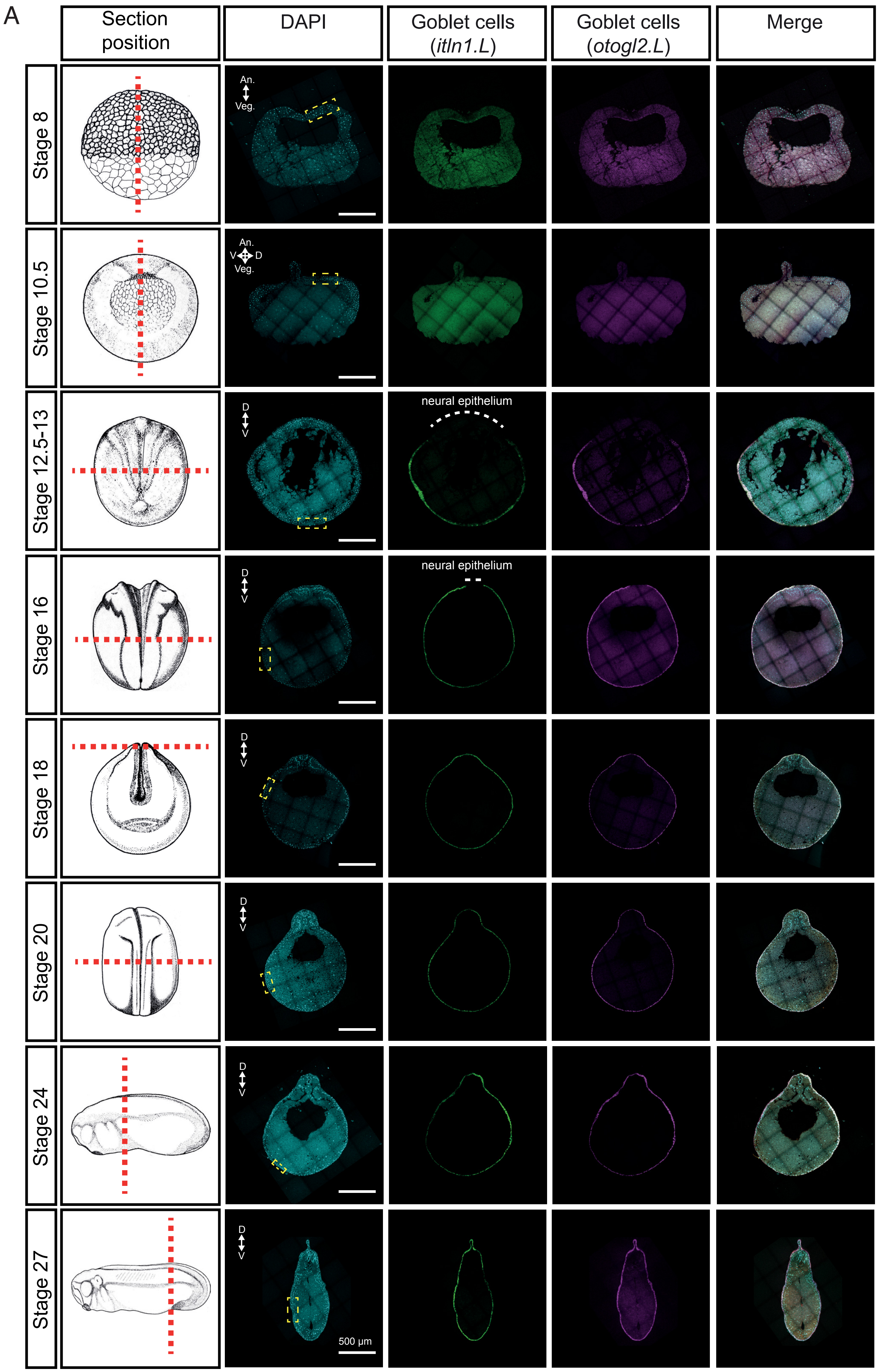

**Fig. S11**(**A**) *In-situ* hybridization chain reaction (HCR) of 8 developmental stages indicating goblet cell differentiation marked by intelectin (*itln1.L*, green) and otogelin (*otog.L/mucXS/otogl2.L*, magenta); nuclei are marked by DAPI staining (cyan). The yellow rectangles indicate regions zoomed-in in Fig. 7B. Images represent maximum intensity projections of transverse sections.

### Supplementary figure 12

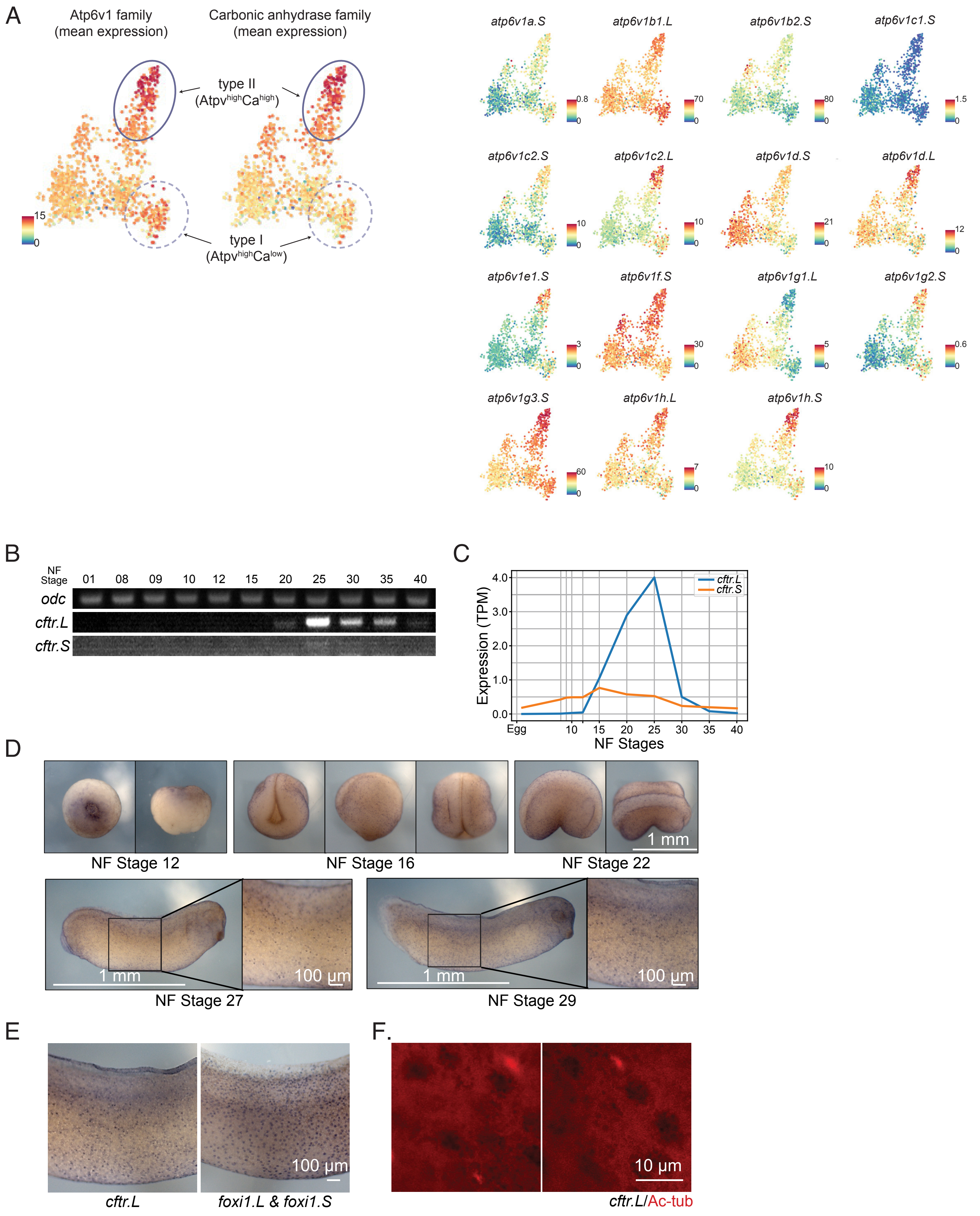

**Fig. S12**(**A**) Low dimensional visualisation of ionocyte subpopulations, overlaid with the mean expression of Atp6v1 and carbonic anhydrase family members (same as Fig. 5D), indicating type-I ($Atp6v1^{high}Ca^{low}$) and type II ionocytes ($Atp6v1^{high}Ca^{high}$). Individual expression of Atp6v1 and carbonic anhydrase family members (right panel).

The scale bars indicate the scaled imputed expression of respective markers.

(**B**) Expression of *cftr.L* and *cftr.S* during MCE developmental stages, indicating peak expression at tailbud stages.

(**C**) Bulk expression from RNA-seq of *cftr.L* and *cftr.S* during MCE differentiation. RNA-seq captures the initial increase in expression during neurula stages and peaking at tailbud stages.

(**D**) Whole-mount *in-situ* staining for *cftr.L* across NF stages 12, 16, 22, 27 and 29, with zoomed-in views of stage 27 and 29

(**E**) Zoomed-in view of whole-mount *in-situ* staining for *cftr.L* and *foxi1.L*-marked ionocytes in tailbud stage embryos.

(**F**) Distinct marking of ciliated cells (red, acetylated-tubulin (Ac-tub)) and ionocytes (black, *cftr.L*) in tailbud stage embryos.

### Supplementary figure 13

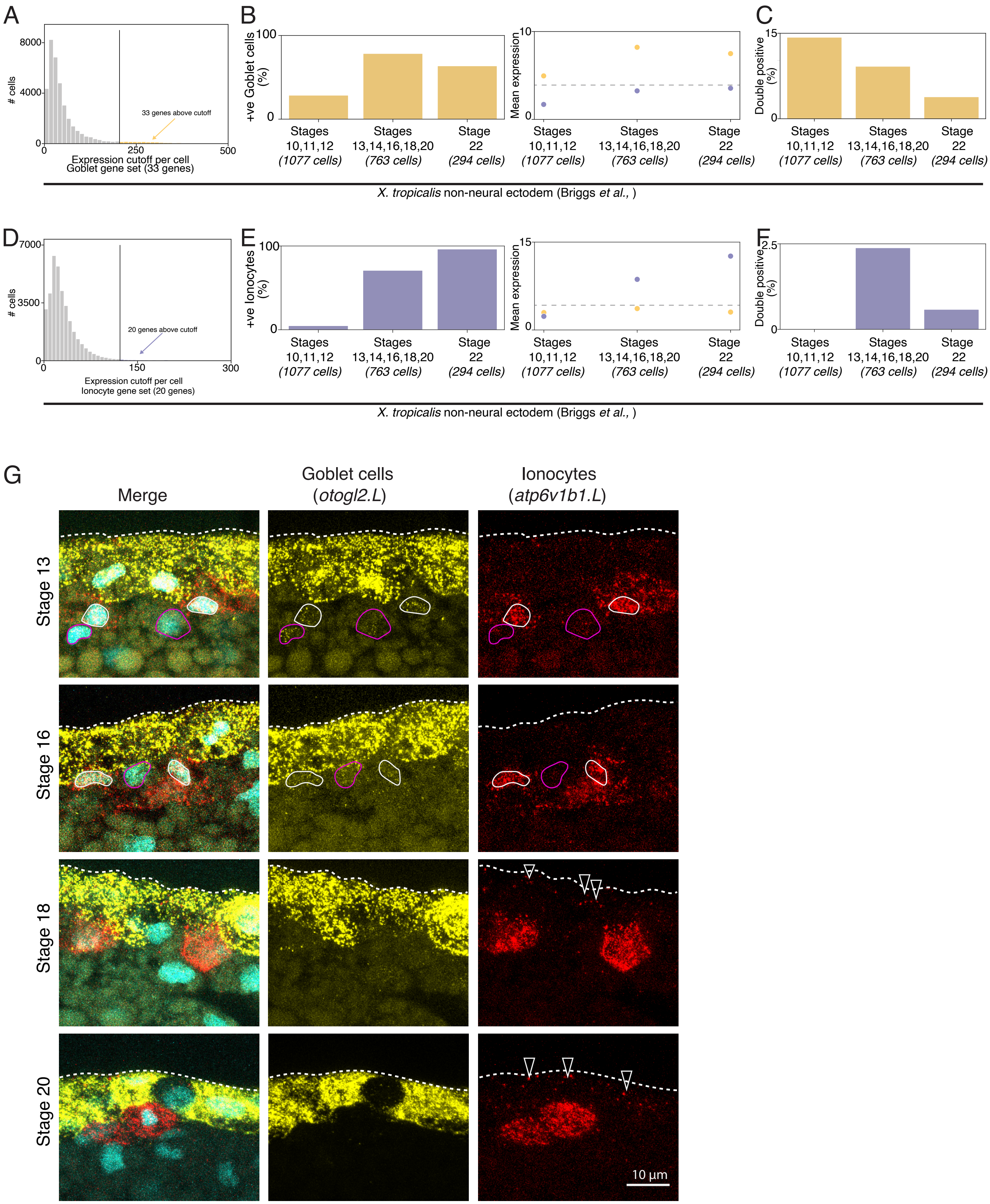

**Fig. S13**(**A**) Expression cutoff to select annotated goblet cells robustly expressing goblet gene signature (33 genes) across *X. tropicalis* development atlas (non-neural ectoderm lineage)

(**B**) The number, percentage and mean expression of goblet cells increases over *X. tropicalis* development atlas (non-neural ectoderm lineage).

(**C**) The number, and percentage of goblet cells expressing ionocyte signature genes (double-positive cells).

(**D**) Similar to (A), but for ionocyte gene signature (20 genes), across *X. tropicalis* development atlas (non-neural ectoderm lineage)

(**E**) Similar to (B), but representing the increase in the number, percentage and mean expression of ionocytes gene signature (20 genes)

(**F**) Similar to (C), but representing the number, percentage of ionocytes expressing goblet cell signature genes (double-positive cells).

(**G**) *In-situ* HCR for goblet cell (*otogl2.L*) and ionocyte marker (*atp6v1b1.L*) across neurula stages. White outlines mark nuclei of double-positive cells within the sensorial layer; pink outlines mark nuclei of cells with low levels of *atp6v1b1.L* and *otogl2.L* expression; white arrowheads mark double-positive cells within the superficial layer. Nuclei are marked by Dapi (cyan). A white dotted line marks the outer boundary of the superficial layer.

Supplementary figure 14

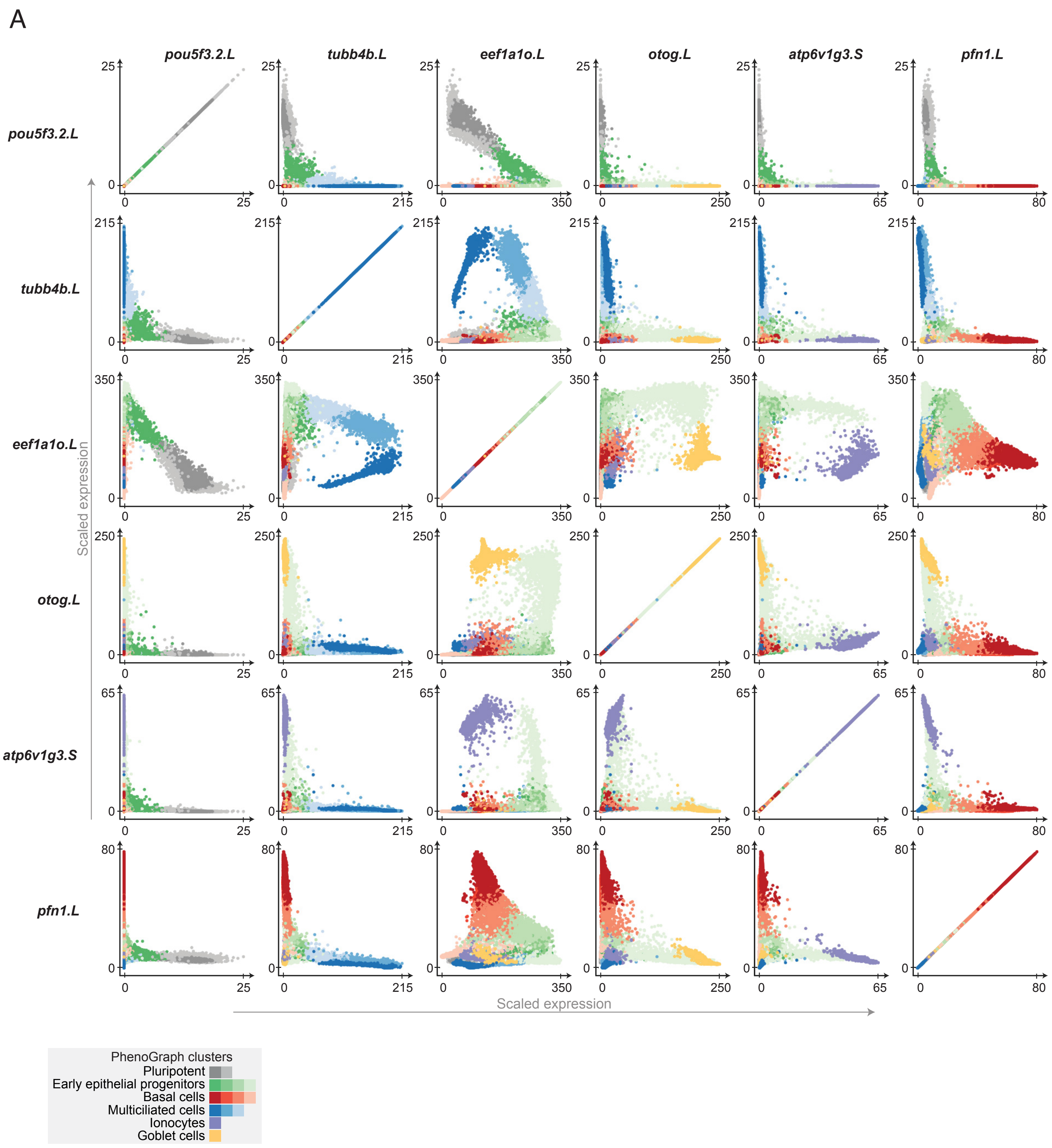

**Fig. S14**(**A**) Cell type marker correlation across individual cells and clusters over MCE development. The early progenitor clusters express multiple cell-types markers at lower levels (notably secretory genes), indicating multi-lineage bias.

### Supplementary figure 15

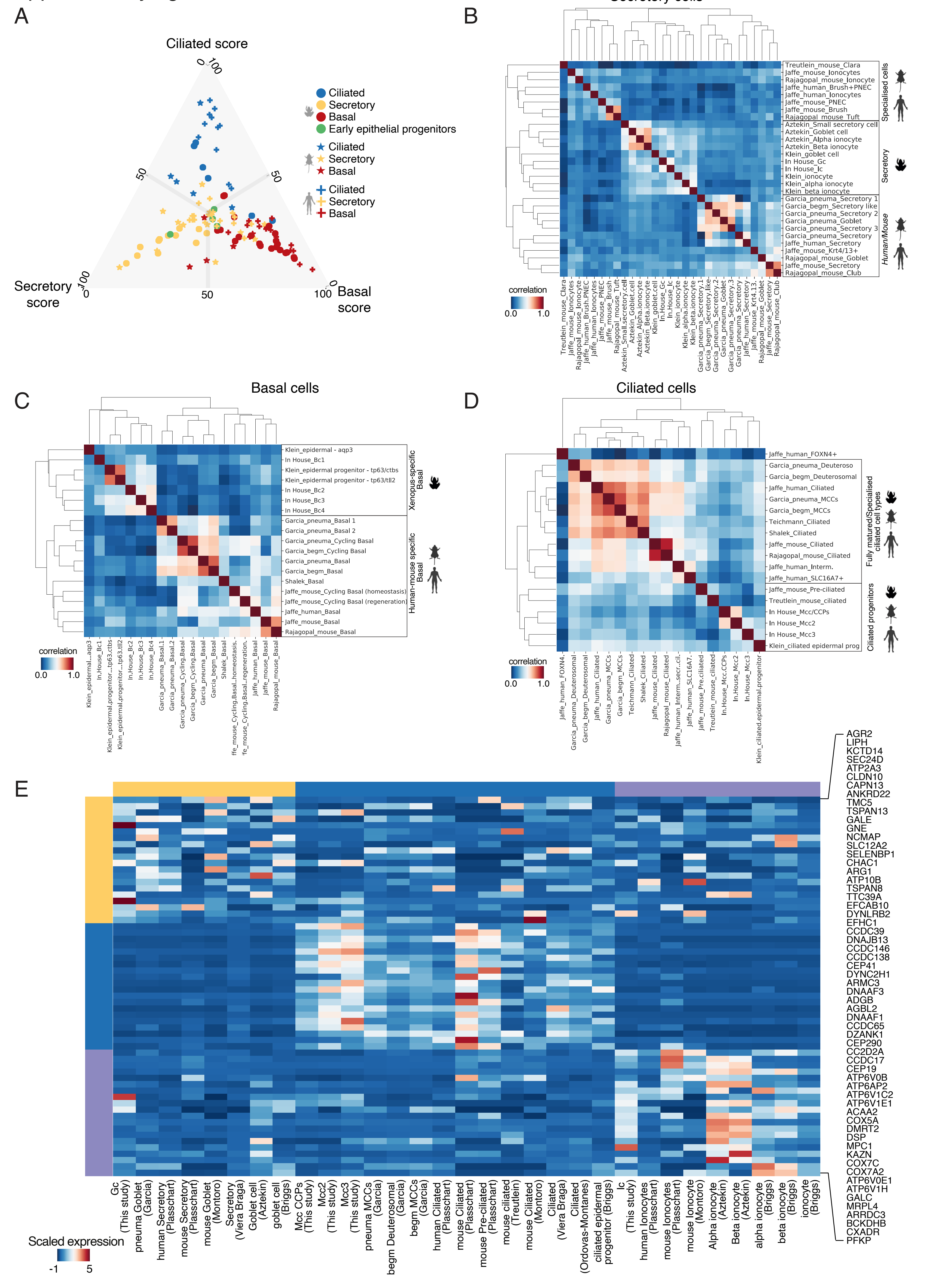

**Fig. S15**(**A**) Classification and scoring of the different individual cell types from *Xenopus*, mouse and human single-cell studies, based on expressed and orthologous gene set. We plot the ciliated, basal and secretory enrichment scores (Fig. 10A) to annotate cell type specificity and accuracy to score respective cell types. The colors indicate the different cell types, while shapes indicate the different species.

(**B**) Correlation of secretory cell types from *Xenopus*, mouse and human single-cell datasets. The *Xenopus* secretory cell types are grouped separately from their higher vertebrate counterparts, likely due to a lack of mucins and other secretory molecules. The specialised secretory cells from higher vertebrates also form a separate cluster.

(**C**) Correlation of basal cell types from *Xenopus*, mouse and human single-cell datasets. The *Xenopus* basal cell types are grouped separately from the higher vertebrate basal cell subtypes, owing to a distinct mode of specification and specialised function across higher vertebrate basal cell types

(**D**) Correlation of ciliated cell types from *Xenopus*, mouse and human single-cell datasets. The ciliated progenitors (irrespective of species) are grouped together between vertebrates and distinctly separated from mature ciliated cell types, indicating a conserved expression module driving ciliogenesis.

(**E**) Marker gene expression (human ortholog DE genes, y-axis) in the different MCE cell types across species (x-axis). Cell type annotations are highlighted in groups (yellow, blue and purple bars across rows and columns). The scale bars indicate the z-scaled mean expression of respective markers.
